# Neuroprotective mechanisms of red clover and soy isoflavones in Parkinson’s disease models

**DOI:** 10.1101/2020.12.01.391268

**Authors:** Aurélie de Rus Jacquet, Abeje Ambaw, Mitali Arun Tambe, Sin Ying Ma, Michael Timmers, Mary H. Grace, Qing-Li Wu, James E. Simon, George P. McCabe, Mary Ann Lila, Riyi Shi, Jean-Christophe Rochet

**Affiliations:** Department of Medicinal Chemistry and Molecular Pharmacology, Purdue University, West Lafayette, IN, 47907, USA; Centre de Recherche du CHU de Québec, Axe Neurosciences, Québec, QC, G1V 4G2, Canada, and Département de Psychiatrie & Neurosciences, Université Laval, Québec, QC, G1V 0A6, Canada; Department of Basic Medical Sciences, Purdue University, West Lafayette, IN, 47907, USA; Physiology Department, Monterey Peninsula College, Monterey, CA, 93940, USA; National Center for Advancing Translational Sciences, National Institutes of Health, Bethesda, MD, USA; Plants for Human Health Institute, Department of Food Bioprocessing and Nutrition Sciences, North Carolina State University, Kannapolis, NC, 28081, USA; Berry Blue, Kannapolis, NC, 28081, USA; Department of Plant Biology, Rutgers University, New Brunswick, NJ, 08901, USA; Department of Statistics, Purdue University, West Lafayette, IN 47907, USA; Weldon School of Biomedical Engineering, Purdue University, West Lafayette, IN, 47907, USA; Purdue Institute for Integrative Neuroscience, Purdue University, West Lafayette, IN, 47907, USA

**Keywords:** alpha-synuclein, astrocytes, flavonoids, mitochondria, neurodegeneration, neuroprotection, Nrf2, 6-OHDA, polyphenols, rotenone

## Abstract

Parkinson’s disease (PD) is a neurodegenerative disorder characterized by nigrostriatal degeneration and the spreading of aggregated forms of the presynaptic protein α-synuclein (aSyn) throughout the brain. PD patients are currently only treated with symptomatic therapies, and strategies to slow or stop the progressive neurodegeneration underlying the disease’s motor and cognitive symptoms are greatly needed. The time between the first neurobiochemical alterations and the initial presentation of symptoms is thought to span several years, and early neuroprotective dietary interventions could delay the disease onset or slow PD progression. In this study, we characterized the neuroprotective effects of isoflavones, a class of dietary polyphenols found in soy products and in the medicinal plant red clover (*Trifolium pratense*). We found that isoflavone-rich extracts and individual isoflavones rescued the loss of dopaminergic neurons and the shortening of neurites in primary mesencephalic cultures exposed to two PD-related insults, the environmental toxin rotenone and an adenovirus encoding the A53T aSyn mutant. The extracts and individual isoflavones also activated the Nrf2-mediated antioxidant response in astrocytes via a mechanism involving inhibition of the ubiquitin-proteasome system, and they alleviated deficits in mitochondrial respiration. Furthermore, an isoflavone-enriched soy extract reduced motor dysfunction exhibited by rats lesioned with the PD-related neurotoxin 6-OHDA. These findings suggest that plant-derived isoflavones could serve as dietary supplements to delay PD onset in at-risk individuals and mitigate neurodegeneration in the brains of patients.

**Graphical Abstract:** The isoflavone-rich extracts red clover and soy and the individual isoflavones daidzein and equol protect neuronal cultures against environmental and genetic triggers of Parkinson’s disease, and rescue motor deficits in rats exposed to the neurotoxin 6-OHDA.

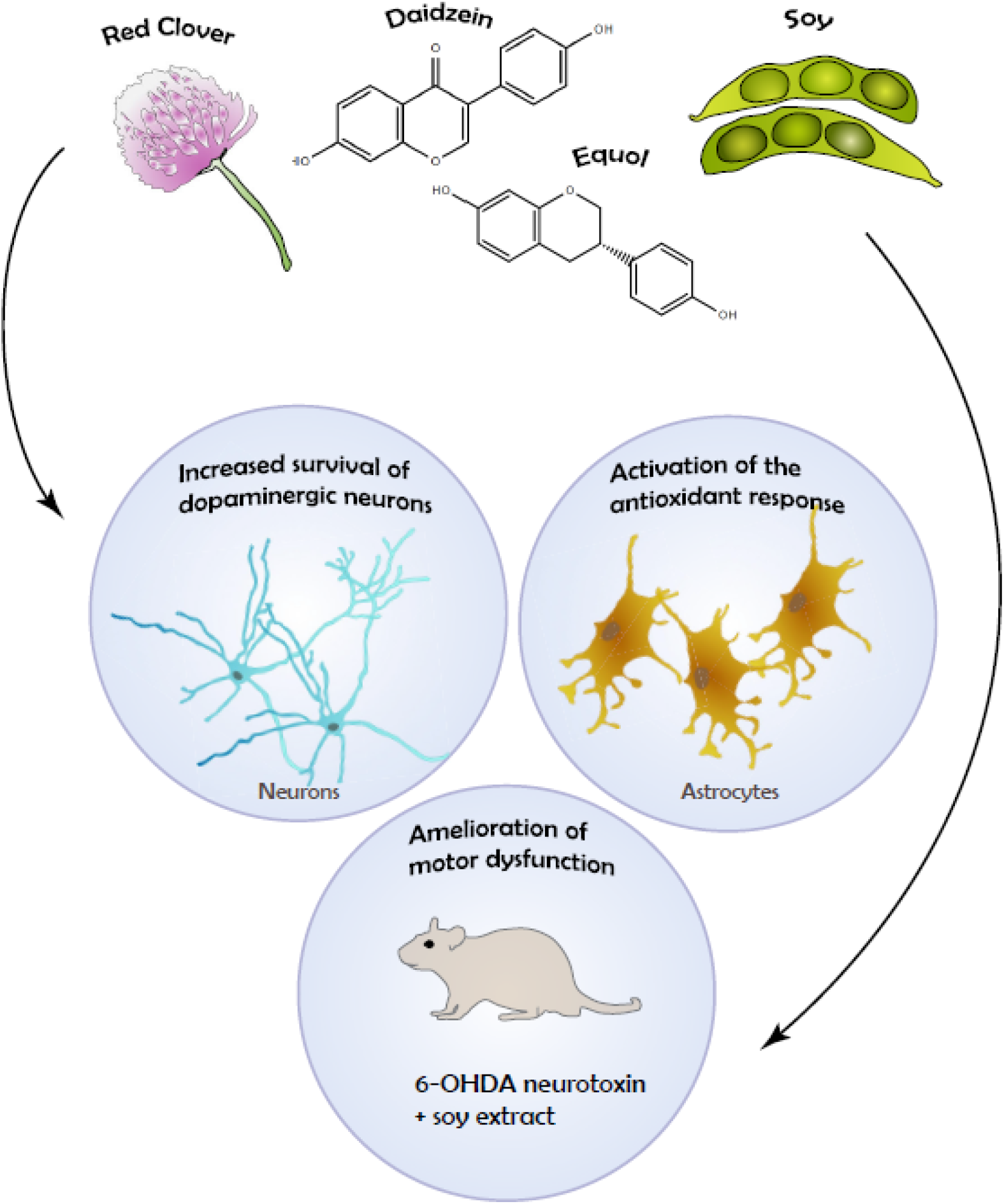

## 1. Introduction

Parkinson’s disease (PD) is a neurodegenerative disorder that affects 5% of the global population over the age of 85^1–3^. The disease involves a loss of dopaminergic neurons from the *substantia nigra* in the midbrain, and this neuronal loss is largely responsible for motor symptoms such as the inability to initiate movement, resting tremor, and reduced balance^4^. Pathological phenomena underlying neurodegeneration in PD include oxidative stress^5, 6^, loss of mitochondrial function^7–9^, aggregation of the presynaptic protein α-synuclein (aSyn)^10–12^, and microglial activation^13^. Familial PD mutations in the *SNCA* gene encoding aSyn are thought to promote the formation of toxic aSyn oligomers by increasing the protein’s expression levels or intrinsic aggregation propensity in the case of duplication/triplication mutations or substitution mutations, respectively^12, 14–18^. Epidemiological evidence suggests that exposure to environmental toxins such as rotenone and paraquat leads to an increase in PD risk^19–22^, potentially via a mechanism involving the production in neurons of reactive oxygen species (ROS) that in turn lead to a build-up of oxidatively modified forms of aSyn with a high propensity to form potentially toxic aggregates^12, 23, 24^. Current PD treatments consist primarily of dopamine replacement therapy (DRT), including administration of the dopamine metabolic precursor L-DOPA or synthetic dopamine receptor agonists, but these agents only alleviate symptoms without delaying the underlying neuronal death^25^. Patients on long-term DRT eventually experience pronounced side effects that include L-DOPA-induced dyskinesias, aggression, and insomnia^26^. Accordingly, there is a critical need to identify disease-modifying therapies that slow PD progression.

Multiple lines of evidence suggest that isoflavone-rich extracts such as soy-derived products and the medicinal plant red clover (*Trifolium pratense*) could have neuroprotective effects in the brains of PD patients. In an ethnopharmacological survey of Pikuni-Blackfeet traditional medicine, we identified red clover as a traditional treatment for symptoms related to PD^27^. Moreover, epidemiological studies revealed that estrogen replacement therapy may be linked to improved cognitive function and delayed onset of PD and Alzheimer’s disease (AD)^28–31^, in turn implying that the consumption of phytoestrogens in isoflavone-rich foods and plant medicines could slow functional decline of the aging brain. In support of this idea, isoflavones and isoflavone-rich extracts were found to exhibit neuroprotective activity in preclinical models of AD and cognitive impairment^32, 33^. Individual isoflavones such as genistein, a major isoflavone found in soy, have also been reported to rescue the PD-like phenotypes of rodents exposed to MPTP or 6-OHDA by restoring motor functions and preserving dopaminergic neurons^34–37^. Although these observations suggest that isoflavones can alleviate neurodegeneration in PD models, the effects of isoflavones or isoflavone-rich botanical extracts on toxicity elicited by insults linked epidemiologically or genetically to PD are poorly understood. Polyphenols (including isoflavones) are well known for their ability to scavenge ROS^38, 39^. However, the fact that brain levels of polyphenols are lower than those of endogenous glutathione suggests that polyphenols may mitigate neurotoxicity via additional protective mechanisms, including the activation of nuclear factor E2-related factor 2 (Nrf2), a transcription factor that regulates the expression of genes involved in the cellular antioxidant response, as well as modulation of the ubiquitin-proteasome system (UPS) and rescue of mitochondrial functional deficits^27, 40^.

This research study was aimed at characterizing isoflavone-rich extracts and individual isoflavones in terms of (i) their neuroprotective effects in cellular models relevant to PD; and (ii) their mechanisms of action, with an emphasis on their ability to stimulate the Nrf2-mediated antioxidant pathway, the UPS, and mitochondrial function. Our findings support the idea that medicinal plants enriched in isoflavones such as red clover and soy could form the basis of dietary interventions for individuals at risk for PD or therapeutic preparations to slow PD progression.

## 2. Materials and methods

### 2.1. Materials

Chemicals were obtained from Sigma Chemical Co. (St. Louis, MO), except when noted. Dulbecco’s Minimal Essential Media (DMEM), fetal bovine serum (FBS), penicillin streptomycin, and trypsin-EDTA were obtained from Invitrogen (Carlsbad, CA). Nuserum was purchased from Thermo Fisher Scientific (Waltham, MA). The SH-SY5Y cells were purchased from ATCC (Manassas, Virginia). Human iPSC-derived astrocytes (iCell astrocytes) were obtained from Cellular Dynamics International (CDI) (Madison, WI). The Novasoy 400 extract was obtained from Archer Daniels Midland (Chicago, IL). The vector pSX2_d44_luc^41^ was provided by Dr. Ning Li (UCLA) with the permission of Dr. Jawed Alam (LSU Health Sciences Center), and the GFPu reporter adenovirus was provided by Dr. Xuejun Wang (University of South Dakota)^42^.

### 2.2. Antibodies

The following antibodies were used in this study: chicken anti-microtubule-associated protein 2 (MAP2) (catalog number CPCA-MAP2, EnCor Biotechnology, Gainesville, FL); rabbit anti-tyrosine hydroxylase (TH) (catalog number 2025-THRAB, PhosphoSolutions, Aurora, CO); anti-rabbit IgG-Alexa Fluor 488 and anti-chicken IgG-Alexa Fluor 594 (Invitrogen, Carlsbad, CA).

### 2.3. Preparation and dissolution of botanical extracts

Red clover flowers were harvested, immediately dried at 37 °C with a food processor, and the water extract was prepared as described previously to reproduce the traditional methods of preparation of red clover-based herbal remedies^27, 40^. The soy extract was prepared as described from Tofu soybeans^43, 44^. Prior to each experiment, extracts were dissolved in sterile deionized water (red clover) or ethanol (soy and Novasoy 400).

### 2.4. HPLC-TOF-MS profiling of a red clover extract

HPLC-TOF-MS analyses were conducted on an Agilent 6220a TOF-MS, equipped with a 1200 series HPLC (Agilent Technologies, Santa Clara, CA) with a Waters X-Bridge column (4.6 x 100 mm, 3.5 μm, Waters, Milford, MA). The separation was conducted using 0.1% (v/v) formic acid in water (solvent A), and 0.1% (v/v) formic acid in acetonitrile (solvent B) with a gradient of 2% B (0-5 min), 2-40% B (5-14 min), 40-100% B (14-17 min), 100% B (17-20 min), and then a return to the original gradient of 2% B (20-24 min). The TOF parameters included a drying gas of 10 L/min, nebulizer pressure of 45 psi, capillary voltage of 3.5 kV, fragment or voltage of 80 kV, and a mass range of *m/z* 100-3000 in both positive and negative modes. The extract was run at 10 mg/mL with an injection volume of 10 μL. The tentative identification of compounds was based on accurate mass measurements (mass error <5 ppm), the generated molecular formula, and information from published work^45, 46^.

### 2.5. Preparation of rat primary cultures

Primary, mixed midbrain cultures were prepared via dissection of E17 embryos obtained from pregnant Sprague-Dawley rats (Harlan, Indianapolis, IN) using methods approved by the Purdue Animal Care and Use Committee^15, 27, 40, 47, 48^. Briefly, the mesencephalic region containing the *substantia nigra* and ventral tegmental area was isolated stereoscopically, and the cells were dissociated with trypsin (final concentration, 26 μg/mL in 0.9% [w/v] NaCl). For experiments that involved analysis of neuroprotective activity (see Section 2.6), the dissociated cells were plated in the wells of a 48- or 96-well, plate (pretreated with poly-L-lysine, 5 μg/mL) at a density of 163,500 or 81,750 cells per well (respectively) in midbrain culture media consisting of DMEM, 10% (v/v) FBS, 10% (v/v) horse serum (HS), penicillin (10 U/mL), and streptomycin (10 μg/mL). After 5 DIV, the cultures were treated with AraC (20 μM, 48 h) to slow the proliferation of glial cells. For experiments that involved monitoring activation of the Nrf2 pathway (see Section 2.7), the dissociated cells were plated in the wells of a 96-well, black clear-bottom plate (pretreated with poly-L-lysine, 10 μg/mL) at a density of 92,650 cells per well in midbrain culture media. After 5 DIV, the cultures were treated with AraC (20 μM, 72 h) before initiating experimental treatments.

Primary midbrain or cortical astrocytes were prepared via dissection of E17 embryos obtained from pregnant Sprague-Dawley rats (Harlan, Indianapolis, IN) using methods approved by the Purdue Animal Care and Use Committee, as described above. The dissociated midbrain cells were plated in a 6-well plate pre-treated with rat collagen (25 μg/mL) at a density of ~270,000 cells per well, while the cortical cells were plated in a 175 cm^2^ flask pre-treated with rat collagen (25 μg/mL) at a density of ~1.45×10^7^ cells per dish. The cultures were maintained in media consisting of DMEM, 10% (v/v) FBS, 10% (v/v) HS, penicillin (10 U/mL), and streptomycin (10 μg/mL). After 48 h, new media consisting of DMEM, 10% (v/v) Nuserum, 10% (v/v) HS, penicillin (20 U/mL), and streptomycin (20 μg/mL) was added to selectively propagate the astrocyte population (clusters of cells that attached the dish) and remove unattached neurons. The media was replaced every two days until most of the astrocytes had spread out on the plate (generally by 7 DIV). The astrocyte-rich culture was passaged at least once (and no more than twice) before being used for the described experiments.

### 2.6. Analysis of neuroprotective activity of red clover or soy extracts in primary midbrain cultures

Neuroprotective activities of botanical extracts and compounds were assessed as described previously^27, 40^. Briefly, primary midbrain cultures (7 DIV) were incubated in the absence or presence of extract or compound for 72 h and then incubated in fresh media containing rotenone (with or without extract or compound) for another 24 h. Alternatively, midbrain cultures were transduced with an adenovirus encoding aSyn A53T (A53T Ad)^15^ at a multiplicity of infection (MOI) of 15 in the absence or presence of extract or compound for 72 h and then incubated in fresh media (with or without extract or compound) for another 24 h. Control cultures were incubated in media without rotenone, A53T Ad, or extract. The cultures were fixed, permeabilized, blocked, and incubated with primary antibodies specific for MAP2 (chicken, 1:2000) and TH (rabbit, 1:1000) for 48 h at 4 °C^15, 27, 40, 47, 48^. The cells were then washed with PBS and incubated with a goat anti-chicken antibody conjugated to Alexa Fluor 594 (1:1000) and a goat anti-rabbit antibody conjugated to Alexa Fluor 488 (1:1000) for 1 h at 22 °C. Relative dopaminergic cell viability was assessed by counting MAP2- and TH-positive neurons in a blinded manner using images taken with an automated Cytation 3 Cell Imaging Reader (BioTek Instruments, Winooski, VT) equipped with a 4X objective. At least 12 images with a total of ~500 to 1000 MAP2^+^ neurons were analyzed per experiment for each treatment, and the data were expressed as the percentage of MAP2^+^ neurons that were also TH^+^ to normalize for variations in cell plating density. Each experiment was performed using at least 3 independent preparations of embryonic cultures.

Neurite length measurements were carried out on images taken with an automated Cytation 3 Cell Imaging Reader equipped with a 4X objective (generally, these images were the same as those used to assess dopaminergic cell viability as outlined above). Lengths of MAP2^+^ processes extending from TH^+^/MAP2^+^ neurons with an intact cell body (~90 neurons per sample) were measured in a blinded manner using the manual length measurement tool of the NIS Elements software (Nikon Instruments, Melville, NY).

### 2.7. Treatment with ARE-EGFP reporter adenovirus

An adenovirus encoding EGFP downstream of the SX2 (E1) enhancer and minimal promoter derived from the mouse heme oxygenase-1 (HO-1) gene^41, 49, 50^ was prepared, and activation of the Nrf2 pathway was monitored in cortical astrocytic cultures and primary midbrain cultures as described^27, 40^. Briefly, primary cortical astrocytes and primary midbrain astrocytes were plated at a density of 5,000 cells per well on a 96-well black clear-bottom plate and transduced with the ARE-EGFP reporter virus at an MOI of 25. Additional experiments were carried out with human iCell astrocytes produced at CDl (Madison, WI) by differentiating an iPSC line that was reprogrammed from fibroblasts obtained from an apparently healthy female individual without known PD-related mutations. Each lot provided by CDI consists of >95% astrocytes that express relevant markers (S100β and GFAP) and respond to pro-inflammatory stimulation by cytokines. iCell astrocytes were plated at a density of 10,000 cells per well on a 96-well, black clear-bottom plate (pretreated with laminin, 10 μg/mL) in DMEM/F12, HEPES media supplemented with 2% (v/v) FBS and 1% (v/v) N-2 supplement. After 24 h, the cells were transduced with the ARE-EGFP reporter virus at an MOI of 25. In other experiments, primary midbrain cultures (8 DIV) prepared as described above (see Section 2.5) were transduced with ARE-EGFP reporter virus at an MOI of 10.

After 48 h, the virus-containing media was removed from the astrocytes or midbrain cultures, and the cells were incubated in fresh media supplemented with botanical extract or compound for 24 h. Control cells were transduced with the ARE-EGFP virus for 48 h and then incubated in fresh media for another 24 h, in the absence of extract (negative control) or in the presence of curcumin (5 μM) (positive control). The cells were incubated in the presence of Hoechst nuclear stain (2 μg/mL in HBSS) for 15 min at 37 °C, washed in HBSS, and imaged in HBSS at 37 °C using a Cytation 3 Cell Imaging Reader equipped with a 4× objective. Quantification of EGFP and Hoechst fluorescence was carried out using the Gen5 2.05 data analysis software (BioTek Instruments). To quantify EGFP fluorescence, regions of interest (ROIs) were generated by the software based on the size range (40 to 400 μm for astrocytes, 20 to 400 μm for mixed midbrain cultures) and a designated fluorescence intensity threshold. For each experiment, the threshold was adjusted so that the overall fluorescence intensity in the curcumin-treated culture was 1.5-to 2.5-fold greater than in the negative-control culture. To quantify Hoechst fluorescence, ROIs were generated by the software based on a size range of 10 to 40 μm. The fluorescence intensity threshold was set so that most of the nuclei stained with Hoechst were included among the detected ROIs. Finally, the number of ROIs for EGFP was divided by the total cell number (ROIs obtained from Hoechst fluorescence) for each treatment and normalized to the control value to obtain a fold-change value.

### 2.8. RNA isolation and quantitative RT-PCR

Cortical astrocytes were harvested by centrifugation at 1,300 x g for 10 min at 4 °C, and the mRNA was extracted using an E.Z.N.A Total RNA kit (Omega Bio-Tek, Norcross, GA). Total RNA (200 ng) was reverse-transcribed using an iScript cDNA Synthesis Kit (Bio-Rad, Hercules, CA). Quantitative PCR (qRT-PCR) was performed using an iQ SYBR Green PCR Kit (Bio-Rad) with forward and reverse primers specific for heme oxygenase 1 (HO1), glutamate-cysteine ligase, catalytic subunit (GCLC), Nrf2, or GAPDH. Primer sequences designed for the rat cDNA sequences were as follows (forward primer sequence relative to the coding sequence listed first for each primer pair): GCLC:^5^’GTTCAACACAGTGGAGGACAA^3^’ and ^5^’GGGACTTAGATGCACCTCCTT^5^’; HO1:^5^’ACAACCCCACCAAGTTCAAA^3^’ and ^5^’CCTCTGGCGAAGAAACTCTG^3^’; Nrf2: ^5^’GAGACGGCCATGACTGATT^3^’ and ^5^’CAGTGAGGGGATCGATGAG^3^’; GAPDH: ^5^’GAACATCATCCCTGCATCCA^3^’ and^5^’CCAGTGAGCTTCCCGTTCA^3^’.

Levels of target gene cDNA (encoding HO1, GCLC, or Nrf2) were normalized to levels of control gene cDNA (encoding GAPDH), and the fold change for samples treated with a red clover extract relative to control samples was calculated using the following formula:

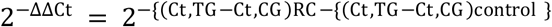

Where Ct,TG represents the crossing threshold for the target gene, Ct,CG represents the crossing threshold for the control gene, ‘RC’ refers to mRNA obtained from cells incubated in the presence of red clover extract, and ‘control’ refers to mRNA obtained from cells incubated in the absence of extract.

### 2.9. UPS reporter assay

UPS function was monitored as described^40^ using an adenoviral reporter construct encoding GFPu, a form of GFP fused C-terminally to the degron CL1^42, 51, 52^. Cortical astrocytes were plated on a 96-well black, clear-bottom plate and transduced with the GFPu reporter adenovirus at an MOI of 2. After 48 h, the cells were incubated in fresh media supplemented with red clover extract for an additional 24 h. Control cells were transduced with the reporter virus for 48 h and then incubated in fresh media for another 24 h, in the absence of extract (negative control) or in the presence of MG132 (2 μM) (positive control). The cells were incubated in the presence of Hoechst nuclear stain (2 μg/mL in HBSS) for 15 min at 37 °C, washed in HBSS, and imaged in HBSS at 37 °C using a Cytation 3 Cell Imaging Reader equipped with a 4X objective.

Quantification of GFP fluorescence was carried out as described previously^40^. Briefly, ROIs were generated by the GEN5 2.05 software based on the cellular size range (20 to 400 μm) and a designated fluorescence intensity threshold. For each experiment, the threshold was adjusted so that a 6-to 7-fold increase in the number of ROIs above the threshold was observed for the MG132-treated culture compared to the negative-control culture. Hoechst fluorescence was quantified by generating ROIs with a fluorescence intensity threshold as described in section 2.7. Finally, the number of ROIs for GFP was divided by the total cell number (ROIs obtained from Hoechst fluorescence) for each treatment and normalized to the control value to obtain a fold-change value.

### 2.10. Oxygen consumption assay

Oxygen consumption was monitored as described previously^40, 53^. Briefly, SH-SY5Y neuroblastoma cells were grown in glucose-free media supplemented with galactose. The cells were incubated in the absence or presence of botanical extract or compound for 19 h and then treated with 30 nM rotenone (with or without extract or compound) for 5 h. Control cells were incubated in the absence of rotenone, extract, or compound for 24 h. The cells were harvested, resuspended in O_2_ consumption buffer (10 mM MgCl_2_, 20 mM HEPES pH 7.2, 8.6 % sucrose (w/v), 0.026 % KH_2_PO_4_ (w/v)), and analyzed for O_2_ consumption rates using a Clark-type oxygen electrode attached to a voltmeter (Digital Model 10 Controller, Rank Brothers, Ltd, Cambridge, UK). An aliquot of 4.5 x 10^6^ cells was loaded into the respiration chamber, and the sample was constantly stirred at 840 rpm. Using the Pico Technology software program (PicoTechnology, Ltd., Cambridgeshire, UK), the O_2_ level remaining in the chamber at any time during respiration was automatically logged (with 10 sec intervals) as VO_2_, the voltage generated by the reaction of O_2_ with the electrode (VO_2_ steadily decreased as O_2_ was consumed). The rate of O_2_ consumption was calculated using the following formula^40, 54^:

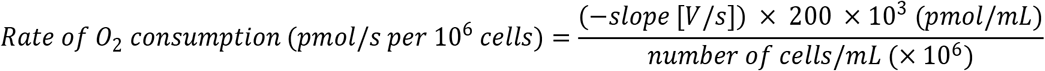

Mean O_2_ consumption rates were determined from 3 independent experiments and normalized to control values.

In other experiments, isoflavone-rich extracts were tested for the ability to displace rotenone from its binding site in complex I of the electron transport chain using a competition assay. Galactose-conditioned SH-SY5Y cells were resuspended in O_2_ consumption buffer containing rotenone (50 nM) with or without red clover extract (1 μg/mL) or soy extract (1 μg/mL). Control cells were resuspended in O_2_ consumption buffer without rotenone or extract. Mean O_2_ consumption rates were determined from 3 independent experiments and normalized to control values as described above.

### 2.11. *In vivo* experiment

A soy extract was examined for protective activity in adult male Sprague-Dawley rats (Harlan, Indianapolis, IN) using methods approved by the Purdue Animal Care and Use Committee. The rats (250 to 300 g) were divided into four groups: control (n = 3), sham (n = 7), 6-OHDA/DMSO (n = 7), and 6-OHDA/soy (n = 7). The control group received no injections. The sham, 6-OHDA/DMSO, and 6-OHDA/soy groups received a stereotaxic injection of saline or 6-OHDA in saline (4 μg/μL). The 6-OHDA/DMSO and 6-OHDA/soy groups received a daily ip injection of 30% (v/v) DMSO vehicle (1 mL/kg) or soy extract (5-20 mg/kg), respectively, over a period of 39 days. Stereotaxic injections were performed unilaterally in the right medial forebrain bundle (MFB) on day 12, with a delivery of 2 μL (1 μL/min), using the coordinates: ML, −1.5 mm; AP, −4.0 mm; DV, −8.5 mm.

Rat motor function was assessed using the open field test 1 d before the injections and at different times from 12 h after surgery to the end of the study^55, 56^. The animals were placed in a Plexiglas activity box (100 × 100 × 20 cm^3^) with food over the center. Eight infrared beams were set in an X-Y matrix, and the position of the rat relative to the beams was monitored at 200 ms intervals with in-house software. The data were used to calculate the percentage of the total box area covered.

### 2.12. Statistical analysis

Data from measurements of primary neuron viability, ARE-EGFP fluorescence, GFPu fluorescence, mitochondrial O_2_ consumption rates, and rat motor function were analyzed via one- or two-way ANOVA with Tukey’s or Dunnett’s multiple comparisons *post hoc* test using GraphPad Prism version 8.0 (La Jolla, CA). Prior to performing these ANOVA analyses, the data were subjected to a square root transformation (neuron viability data) or log transformation (fluorescence, O_2_ consumption rate, and open field test data) to conform to ANOVA assumptions. Neurite length data were subjected to a log transformation to account for skewness in the data. The log-transformed data were analyzed using an approach that accounts for (i) the possibility of multiple neurites arising from a single cell, and (ii) comparison across experiments conducted on different days. Log-transformed neurite lengths for multiple treatment groups were compared using a general linear model implemented in the ‘Mixed’ procedure of SAS Version 9.3 followed by Tukey’s multiple comparisons *post hoc* test (Cary, NC). For measurements of mRNA expression, fold-change values were log-transformed, and the transformed data were analyzed using GraphPad Prism 8.0 via a one-sample t-test to determine whether the mean of the log(fold-change) was different from the hypothetical value of 0 (corresponding to a ratio of 1). The ‘n’ values specified in the figure legends represent the number of biological replicates (i.e. independent experiments involving cultures prepared at different times).

## 3. Results

### 3.1. Study design

The central hypothesis of this study was that botanical extracts rich in isoflavones can alleviate neurotoxicity elicited by insults linked epidemiologically or genetically to PD. To address this hypothesis, we characterized two extracts with high levels of isoflavones – a red clover extract^57^ and a soy extract prepared from Tofu soybeans^43, 44^ – in terms of their ability to interfere with dopaminergic cell death in primary midbrain cultures exposed to two PD stresses: (i) rotenone, an environmental toxicant epidemiologically linked to increased PD risk^22^; and (ii) virus encoding A53T aSyn, a familial mutant form of aSyn that exhibits accelerated fibrillization and enhanced neurotoxicity compared to the wild-type protein^15, 58, 59^. Moreover, we characterized an isoflavone constituent of the soy extract, daidzein, and a second isoflavone, equol (produced from daidzein by the intestinal microbiota), to determine their effects on dopaminergic neuron survival in primary midbrain cultures exposed to rotenone or A53T Ad. The extracts and individual isoflavones were also examined for their effects on Nrf2 signaling and the UPS, as well as their ability to enhance mitochondrial respiration.

### 3.2. Effects of a red clover extract on dopaminergic cell death elicited by PD-related insults in primary midbrain cultures

Our first objective was to measure the neuroprotective activity of a red clover extract in a rat primary midbrain culture model. Our rationale for using this model was that these cultures consist of post-mitotic dopaminergic and GABAergic neurons and glial cells (e.g. astrocytes and microglia), and thus they reproduce key features of the midbrain region affected in PD patients^47, 60^. The red clover extract was prepared from dried flowers, as described previously by Pikuni-Blackfeet traditional healers^27^, and chemical analysis of the extract via HPLC coupled with UV and MS detection revealed an abundance of isoflavones, including formononetin, biochanin A, and pratensin glycosides, and confirmed the isoflavone-rich nature of the red clover extract (Table 1).

**Table 1.** HPLC-TOF-MS analysis for identification of a red clover extract.

| Rt<sup>a</sup><br>(min) | Experimental<br>*m/z*[M+H]<sup>+</sup> | Theoretical<br>*m/z* [M+H]<sup>+</sup> | Experimental<br>*m/z* [M-H]<sup>-</sup> | Theoretical<br>*m/z* [M-H]<sup>-</sup> | Formula | Tentative identification<sup>b</sup> |
| --- | --- | --- | --- | --- | --- | --- |
| 3.38 | - | - | 191.0188 | 191.0192 | C<sub>6</sub>H<sub>8</sub>O<sub>7</sub> | citric acid |
| 10.27 | 166.0858 | 166.0865 | - | - | C<sub>9</sub>H<sub>11</sub>NO<sub>2</sub> | phenylalanine |
| 12.11 | 205.0967 | 205.0977 | - | - | C<sub>11</sub>H<sub>12</sub>N<sub>2</sub>O<sub>2</sub> | tryptophan |
| 13.60 | 360.1079 | 360.1083 | 358.0921 | 358.0927 | C<sub>18</sub>H<sub>17</sub>NO<sub>7</sub> | cis and trans clovamide |
| 13.93 | 360.1075 | 360.1083 | 358.0935 | 358.0927 | C<sub>18</sub>H<sub>17</sub>NO<sub>7</sub> | cis and trans clovamide |
| 14.45 | 465.1033 | 465.1033 | 463.0894 | 463.0877 | C<sub>21</sub>H<sub>20</sub>O<sub>12</sub> | quercetin-3-O-β-glucoside |
| 14.82 | 551.1033 | 551.1037 | 549.0900 | 549.0881 | C<sub>24</sub>H<sub>22</sub>O<sub>15</sub> | quercetin-3-O-β-malonyl-glucoside |
| 14.90 | 449.1086 | 449.1084 | - | - | C<sub>21</sub>H<sub>20</sub>O<sub>11</sub> | unknown flavonoid glycoside |
| 15.55 | 565.1195 | 565.1193 | - | - | C<sub>25</sub>H<sub>24</sub>O<sub>15</sub> | 3-methylquercetin-7-O-β-glucoside malonate |
| 16.33 | 549.1244 | 549.1244 | - | - | C<sub>25</sub>H<sub>24</sub>O<sub>14</sub> | pratensein-7-O-β-glucoside-6"-O-malonate |
| 17.85 | 547.1089 | 547.1088 | - | - | C<sub>25</sub>H<sub>22</sub>O<sub>14</sub> | irilone-4'-O-β-glucoside-6"-O-malonate |
| 18.14 | 533.1292 | 533.1295 | 1063.2379<br>[2M-H]<sup>-</sup> | 1063.2355 | C<sub>25</sub>H<sub>24</sub>O<sub>13</sub> | biochanin A-7-O-β-glucoside-6"-O-malonate |
| 18.71 | 269.0813 | 269.0814 | 267.0660 | 267.0657 | C<sub>16</sub>H<sub>12</sub>O<sub>4</sub> | formononetin |

<sup>a</sup>Rt: retention time in minutes; <sup>b</sup>compounds were tentatively identified based on accurate mass measurements (with mass error <5 ppm), the generated molecular formula, retention time values, and information from the literature <sup>45, 46</sup>.

In one set of experiments, primary midbrain cultures were exposed to rotenone in the absence or presence of the red clover extract. The cultures were co-stained for MAP2, a general neuronal marker, and TH, a selective marker of dopaminergic neurons (Fig. 1A), and scored for relative dopaminergic neuron viability by determining the ratio of TH^+^ neurons to total MAP2^+^ neurons^15, 27, 40, 47^. Cultures exposed to rotenone alone exhibited a ~20-30% decrease in relative dopaminergic cell viability, and this decrease was abrogated in cultures exposed to rotenone plus red clover extract (Fig. 1B). Moreover, neurite length measurements revealed a decrease in the lengths of processes extending from MAP2^+^/TH^+^ neurons in cultures exposed to rotenone compared to control cultures, consistent with the ‘dying back’ degeneration of dopaminergic neurites that is thought to precede cell death observed in the brains of PD patients^61, 62^, and this decrease in neurite lengths was alleviated in cultures treated with rotenone in the presence of red clover extract (Fig. 1C). These results suggested that polyphenols in the red clover extract interfered with dopaminergic cell death and neurite retraction elicited by rotenone.

**Figure 1.**
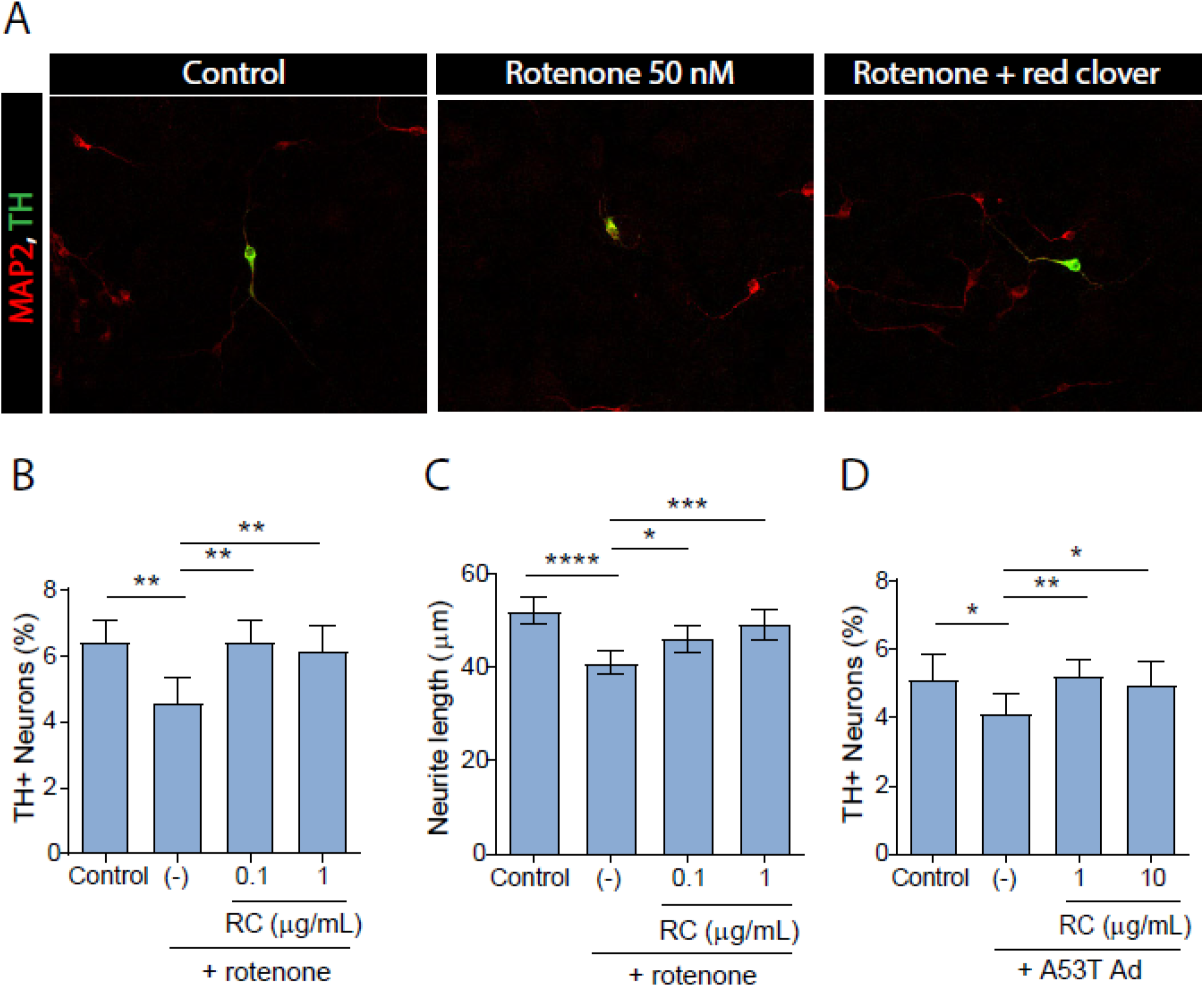
A red clover extract alleviates rotenone and aSyn neurotoxicity. (A) Confocal images of immunostained neurons in primary midbrain cultures show expression of the pan-neuronal marker MAP2 (red) and the dopaminergic neuron marker TH (green). Images show a protective effect of red clover extract against rotenone-induced neurite retraction. (B-D) Cultures were treated with 50 nM rotenone for 24 h (B, C) or A53T Ad (MOI 15) for 72 h (D) in the absence or presence of red clover extract (‘RC’). Control cells were incubated in the absence of rotenone, A53T Ad, or extract. The cells were stained with antibodies specific for MAP2 and TH and scored for relative dopaminergic cell viability (B, D) or neurite length (C). Cell viability data are presented as the mean ± SEM; n = 3; *p≤0.05, **p≤0.01, square root transformation, one-way ANOVA with Tukey’s multiple comparisons *post hoc* test. Neurite length data are presented as the mean value ± 95% confidence limits after back-transformation of log-transformed data as outlined in the Methods; n = 3; *p≤0.05, ***p≤0.001, ****p≤0.0001, Tukey’s multiple comparisons *post hoc* test after log transformation and general linear model implementation.

In a second set of experiments, primary midbrain cultures were transduced with A53T Ad, previously shown to be selectively toxic to MAP2^+^/TH^+^ neurons^15^, in the absence or presence of red clover extract. The cultures were co-stained for TH and MAP2 and analyzed for relative dopaminergic cell survival as outlined above. Cultures treated with virus plus extract had a higher percentage of TH^+^ neurons compared to cultures treated with virus alone (Fig. 1D), suggesting that red clover polyphenols inhibited aSyn-mediated dopaminergic neurotoxicity.

### 3.3. Effects of a red clover extract on astrocytic Nrf2/ARE signaling

The transcription factor Nrf2 is a master regulator of the cellular antioxidant response^63^. In the cytoplasm, Nrf2 is sequestered by Kelch-like ECH-associated protein 1 (Keap1)^64^ and targeted for degradation by the UPS^65^. Under conditions of oxidative stress, the interaction between Nrf2 and Keap1 is disrupted, resulting in Nrf2 translocation to the nucleus where it binds AREs in the regulatory region of its target genes^66, 67^. The Nrf2-mediated response is primarily activated in astrocytes, enabling their production and secretion of glutathione metabolites that are subsequently taken up by neighboring neurons^68, 69^.

Evidence suggests that rotenone and aSyn elicit neurotoxicity at least in part by triggering an increase in oxidative stress^9, 70, 71^. Thus, we hypothesized that the red clover extract could protect against neurotoxicity elicited by rotenone or A53T aSyn by activating the Nrf2-mediated antioxidant response^72^. To address this hypothesis, primary cortical astrocytes were transduced with a reporter adenovirus encoding EGFP downstream of DNA regulatory sequences encompassing two antioxidant response elements (AREs) and a minimal promoter derived from the mouse HO-1 gene^27, 40, 41, 49, 50^. The transduced cultures were incubated in the absence or presence of red clover extract for 24 h and examined for levels of EGFP fluorescence (Fig. 2A). Astrocytes treated with extract exhibited greater fluorescence compared to untreated cells, suggesting that red clover polyphenols activated the Nrf2/ARE antioxidant response (Fig. 2B-C). We observed a similar effect in mixed primary cortical cultures consisting of neurons, astrocytes, and microglia, suggesting that the presence of neurons and other glial cell types did not affect the ability of the red clover extract to activate Nrf2 transcriptional activity in astrocytes (Fig. 2D). To confirm that the observed increase in EGFP fluorescence reflected an increase in Nrf2-mediated transcription, we quantified mRNAs encoding Nrf2 target genes by qRT-PCR. In agreement with the results obtained from the EGFP reporter assay, we found that astrocytes treated with red clover extract exhibited an up-regulation of mRNA encoding HO1 or GCLC compared to control cells (Fig. 2E, F), and the observed timing of expression of these two Nrf2-regulated genes was consistent with previous findings^73^.

**Figure 2.**
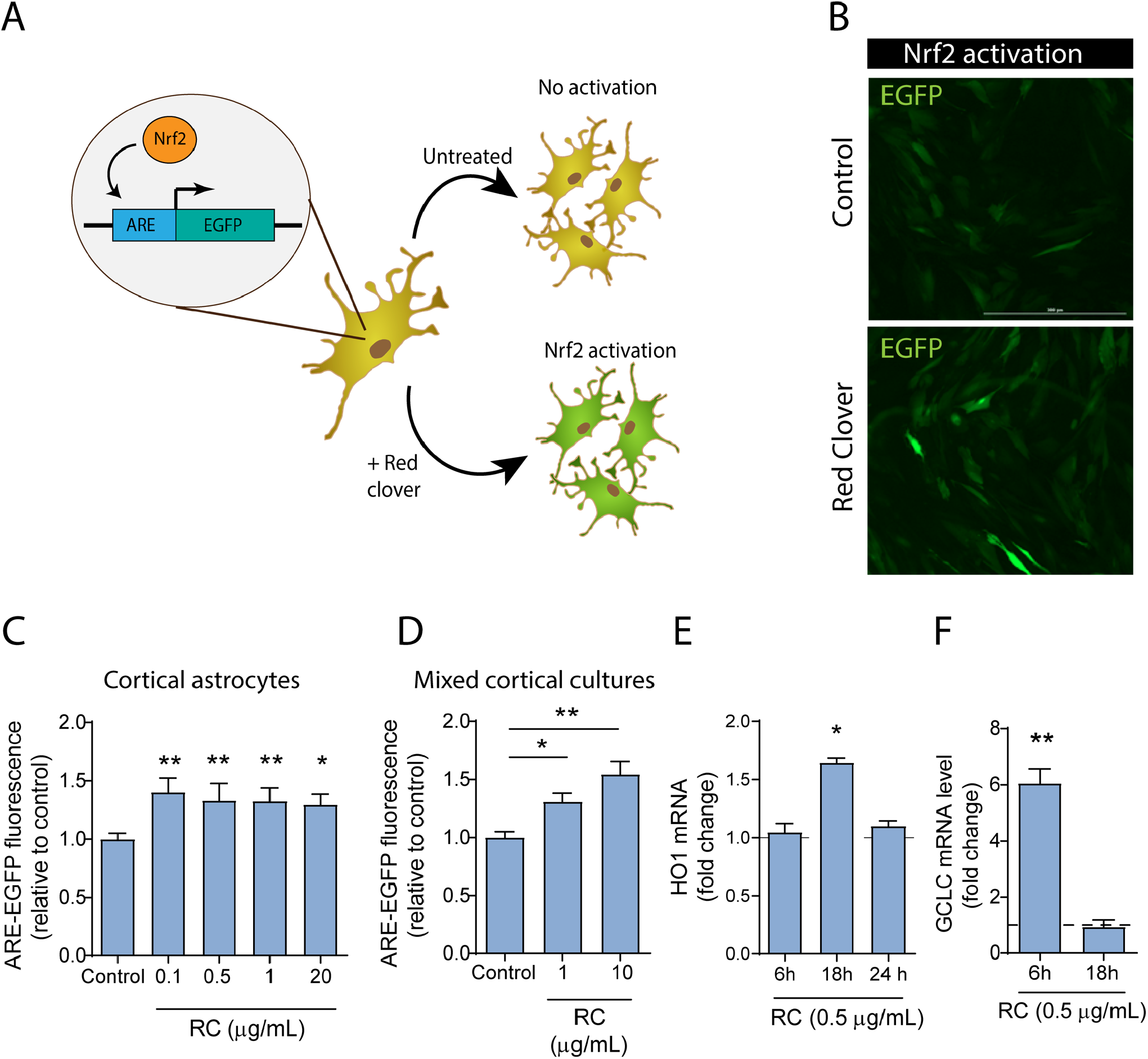
A red clover extract activates the Nrf2/ARE antioxidant response in cortical astrocytes. (A-B) Primary cortical astrocytes were transduced with an ARE-EGFP reporter adenovirus for 48 h and incubated in the absence (‘control’) or presence of red clover extract for 24 h, as illustrated in the schematic (A). The cells were imaged using a Cytation 3 Cell Imaging Reader (scale bar, 300 μm) (B). (C-D) Cortical astrocytes (C) and mixed cortical cultures (D) transduced with ARE-EGFP reporter virus were incubated in the absence (‘control’) or presence of red clover extract (‘RC’) and imaged to determine the intracellular EGFP fluorescence intensity. (E-F) Cortical astrocytes were incubated in the absence or presence of red clover extract, and levels of mRNA encoding HO1 (E) or GCLC (F) were measured by qRT-PCR. The data are presented as the mean ± SEM; n = 4 (C), n = 3 (D,F), n = 2 (E); *p≤0.05, **p≤0.01 versus control, log transformation, oneway ANOVA with Dunnett’s multiple comparison *post hoc* test (C,D); *p≤0.05, **p≤0.01 versus a predicted ratio of 1, log transformation followed by one-sample t-test (E,F).

In subsequent experiments, we assessed whether the red clover extract could activate the Nrf2/ARE pathway in the same cellular context as that in which the extract conferred neuroprotection against toxicity elicited by rotenone or A53T aSyn. To address this question, primary midbrain cultures or astrocytes isolated from these cultures were transduced with the ARE-EGFP reporter adenovirus and incubated in the absence or presence of extract. Surprisingly, isolated astrocytes and neuron-glia cultures from embryonic rat midbrain exhibited similar levels of EGFP fluorescence after incubation with or without extract (Supplementary Fig. 1A, B). In summary, our data indicate that the ability of a red clover extract to activate Nrf2 signaling in cultured astrocytes depends on the brain region from which they were derived (e.g. cortex or midbrain), and mechanisms of Nrf2 activation mediated by the red clover extract and reflected by our ARE-EGFP reporter may not be operative in midbrain cultures.

### 3.4. Mechanism of red clover-mediated Nrf2 activation in cortical astrocytes

Polyphenols have been reported to up-regulate Nrf2 transcriptional activity via various mechanisms. For example, they can trigger Keap1 modification, either by reacting directly with Keap1 regulatory cysteine residues or indirectly by promoting ROS-mediated Keap1 oxidation^74–78^. In turn, Keap1 modification results in disruption of the Nrf2-Keap1 interaction, Nrf2 stabilization, and an increase in Nrf2 nuclear translocation and transcriptional activity. Some polyphenols interfere with the UPS^79–83^, and a loss of UPS function has been linked to the activation of Nrf2 and an increase in the expression of its target genes^84–86^. Other polyphenols, including the flavonoid curcumin, have also been shown to induce increased expression of the Nrf2 gene^87–89^. In the next phase of our study, we carried out experiments aimed at elucidating which of these mechanisms could be involved in the activation of Nrf2 signaling in cortical astrocytes by the red clover extract.

To determine whether a pro-oxidant effect was necessary for Nrf2 activation by the red clover extract, we tested the effect of the extract on Nrf2 transcriptional activity in primary cortical astrocytes cultured in the absence or presence of N-acetyl cysteine (NAC), a cell-permeable antioxidant molecule that serves as a ROS scavenger and a glutathione precursor^48, 60^. Astrocytes transduced with the ARE/EGFP reporter virus and treated with red clover extract plus NAC exhibited similar fluorescence levels compared to transduced astrocytes treated with extract alone (Fig. 3), suggesting that the activation of Nrf2 transcriptional activity by the extract was not affected by treatment with NAC, and, therefore, occurred via a ROS-independent mechanism^48, 60^.

**Figure 3.**
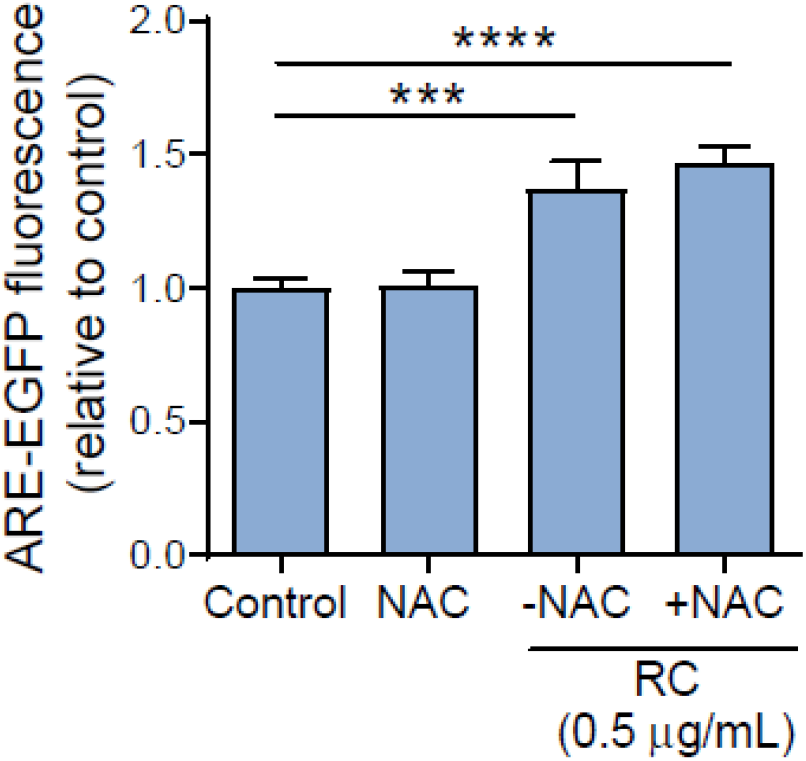
Red clover-mediated activation of the Nrf2/ARE antioxidant response is ROS-independent. Primary cortical astrocytes were transduced with an ARE-EGFP reporter adenovirus for 48 h and incubated with or without a red clover extract (‘RC’) for 24 h in the absence or presence of NAC (1 mM). Control astrocytes were transduced with the reporter virus and incubated in the absence of extract or NAC. The cells were imaged to determine the intracellular EGFP fluorescence intensity. The data are presented as the mean ± SEM; n = 6; ***p≤0.001, ****p≤0.0001, log transformation, one-way ANOVA with Dunnett’s multiple comparisons *post hoc* test.

To address whether Nrf2 activation by the red clover extract could involve UPS inhibition, we tested the effect of the extract on levels of GFP fluorescence in primary cortical astrocytes transduced with a reporter adenovirus encoding GFPu, an unstable form of GFP that is linked to the CL1 degron^40, 42, 51, 52^. Under physiological conditions, GFPu is efficiently degraded, whereas inhibition of the UPS results in accumulation of GFPu and increased cellular fluorescence. GFPu-expressing astrocytes exhibited greater fluorescence when cultured in the presence versus the absence of the red clover extract (Fig. 4A), suggesting that polyphenols in the extract inhibit UPS activity. As a control, we confirmed that UPS inhibition led to the activation of Nrf2 signaling in our cortical astrocyte model by showing that astrocytes transduced with the ARE-EGFP reporter virus showed an increase in EGFP fluorescence when cultured in the presence of the proteasome inhibitor MG132 (Fig. 4B). Curcumin, a polyphenol previously reported to interfere with UPS function^90^, induced an increase in GFP fluorescence in GFPu-expressing astrocytes (Supplementary Fig. 2A) and potently activated the Nrf2 pathway in rat cortical astrocytes, rat mixed cortical or midbrain cultures, and human iPSC-derived astrocytes (Supplementary Fig. 2B), further supporting a link between UPS inhibition and Nrf2 activation.

**Figure 4.**
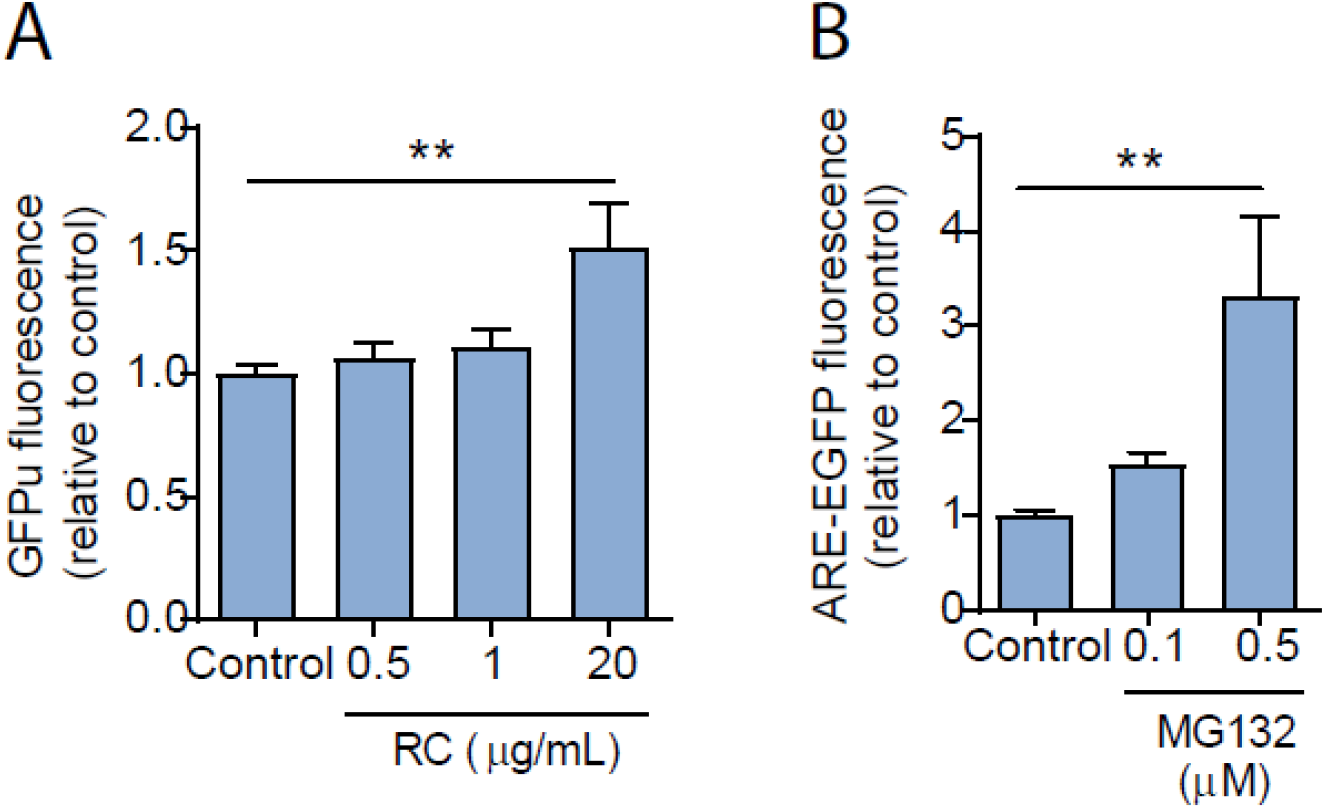
Red clover polyphenols inhibit the UPS. (A) Primary cortical astrocytes transduced with a reporter adenovirus encoding the UPS substrate GFPu for 48 h were incubated in the absence (‘control’) or presence of red clover extract (‘RC’) for 24 h and imaged to determine the intracellular GFP fluorescence intensity. (B) Primary cortical astrocytes transduced with an ARE-EGFP reporter adenovirus for 48 h were incubated in the absence (‘control’) or presence of the proteasome inhibitor MG132 for 24 h and imaged to determine the intracellular EGFP fluorescence intensity. The data in (A) and (B) are presented as the mean ± SEM; n = 3 or 4; **p≤0.01, log transformation, one-way ANOVA with Dunnett’s multiple comparisons *post hoc* test.

To assess whether Nrf2 activation by the red clover extract could involve an increase in the expression of the Nrf2 gene, we quantified Nrf2 mRNA in cortical astrocytes via qRT-PCR. The results indicated that Nrf2 mRNA levels were increased by ~15% in cells incubated with extract for 24 h compared to untreated cells, with a similar trend evident in cells treated with extract for 6 h (Supplementary Fig. 3).

Collectively, these data suggest that polyphenols in the red clover extract inhibit UPS activity and induce a modest increase in Nrf2 mRNA levels in cortical astrocytes. We infer that both of these effects could account at least in part for the ability of the red clover extract to activate Nrf2 signaling in these cells.

### 3.5. Neuroprotective effects of a soy extract and individual isoflavones

Our next objective was to examine the neuroprotective activities of an isoflavone-rich soy extract and two individual isoflavones, daidzein and equol (Fig. 5A), against toxicity elicited by PD-related insults in primary midbrain cultures. Previous HPLC analyses of the phytochemical composition of soy extracts have revealed an enrichment in isoflavones including genistein and daidzein (genistein/daidzein ratio consistently around 1.4), and lower levels of glycitein^43, 44^. Our rationale for characterizing effects of daidzein is that it is an important constituent of soy products, and equol was examined because it is a metabolic product of daidzein in the intestine (i.e. it has translational significance) and exhibits greater antioxidant activity than daidzein itself^91–93^. Cultures treated with rotenone or A53T Ad plus soy extract showed a strong trend towards greater dopaminergic cell viability compared to cultures treated with rotenone or A53T virus alone (Fig. 5B,F). Furthermore, rotenone neurotoxicity was alleviated in midbrain cultures treated with daidzein (100 nM, with a trend towards a neuroprotective effect at 50 nM) or equol (50 nM) (Fig. 5C-E). An increase in dopaminergic neuron viability was also observed in A53T-expressing cultures treated with daidzein at a concentration as low as 25 nM (Fig. 5G). These data suggest that soy polyphenols and the isoflavones daidzein and equol interfere with neurotoxicity elicited by rotenone and A53T aSyn.

**Figure 5.**
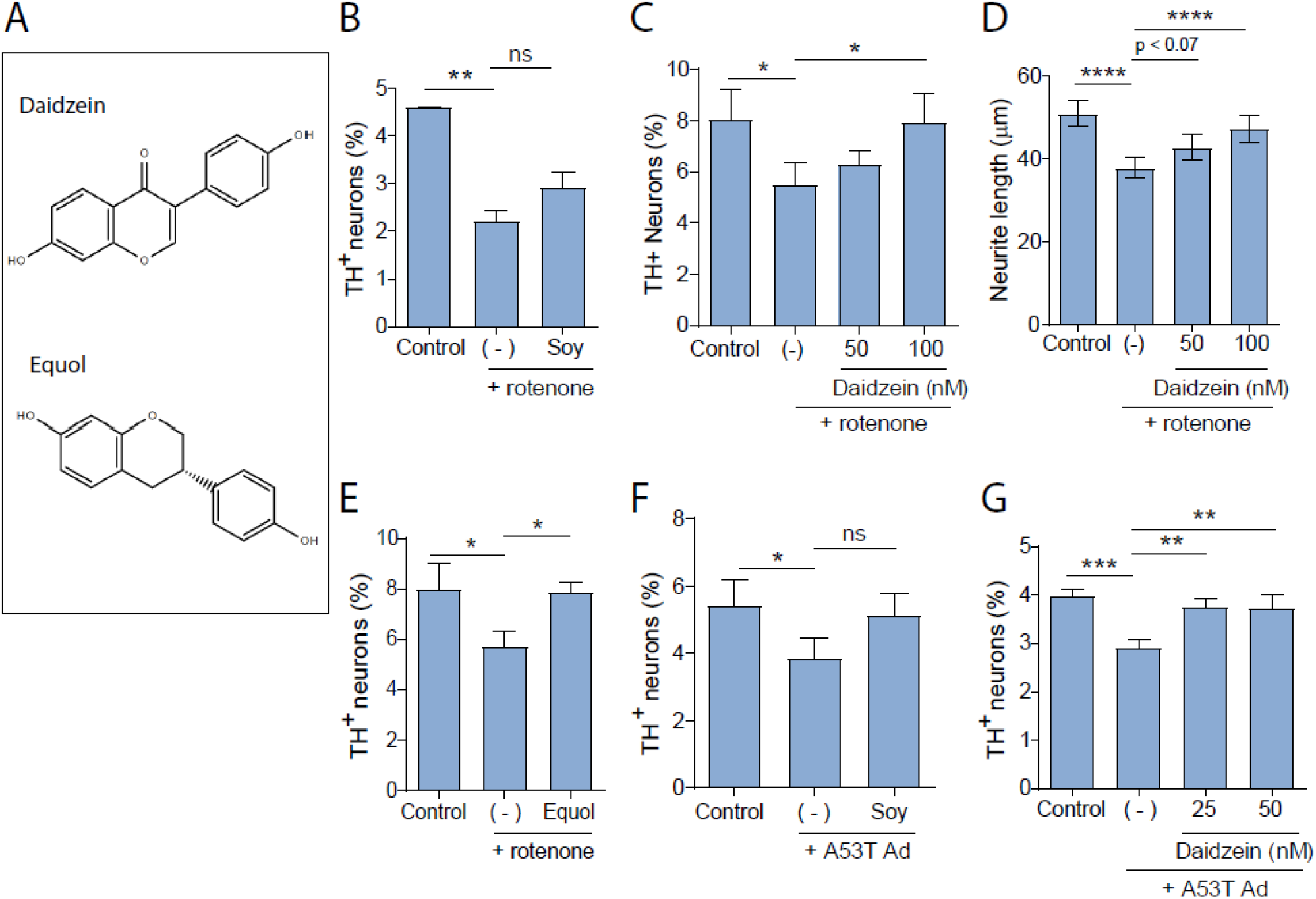
Soy isoflavones alleviate rotenone and aSyn neurotoxicity. (A) Molecular structures of the isoflavones daidzein and equol. (B-G) Primary midbrain cultures were treated with rotenone (50 nM or 100 nM) for 24 h (B-E) or A53T Ad (MOI 15) for 72 h (F,G) in the absence or presence of soy extract (1 μg/mL, B, F), daidzein (25 to 100 nM, C, D, G), or equol (50 nM, E). Control cells were incubated in the absence of rotenone, A53T Ad, or extract/compound. The cells were stained with antibodies specific for MAP2 and TH and scored for relative dopaminergic cell viability (B, C, EG) or neurite length (D). Cell viability data are presented as the mean ± SEM; n = 3 (B, E-G) or n = 4 (C); *p≤0.05, **p≤0.01, ***p≤0.001, square root transformation, one-way ANOVA with Tukey’s multiple comparisons *post hoc* test. Neurite length data (D) are presented as the mean value ± 95% confidence limits after back-transformation of log-transformed data as outlined in the Methods (n = 3); ****p≤0.0001, Tukey’s multiple comparisons *post hoc* test after log transformation and general linear model implementation. In panels (B) and (F), a statistically significant neuroprotective effect is observed for cultures treated with soy extract plus insult versus insult alone when the square root-transformed data are analyzed via ANOVA with the Newman-Keuls *post hoc* test (p≤ 0.05).

In the next set of experiments, we examined whether the soy extract, daidzein, or equol could activate the astrocytic Nrf2/ARE antioxidant response. To address this question, primary cortical astrocytes were transduced with the ARE/EGFP reporter adenovirus and incubated in the absence or presence of the extract or individual isoflavones. Quantification of EGFP fluorescence revealed that none of the treatments induced an increase in the EGFP signal (Supplementary Fig. 4). However, in contrast to a red clover extract, the soy extract activated the Nrf2 response in human iPSC-derived astrocytes (Fig. 6). Additional analyses revealed that cortical astrocytes transduced with the GFPu reporter virus exhibited a modest increase in GFP fluorescence when treated with the soy extract (at a relatively high concentration of 20 μg/mL) or equol, but no increase in fluorescence upon incubation with daidzein (Supplementary Fig. 5). These data suggest that (i) the neuroprotection elicited by soy polyphenols, daidzein, and equol may not rely on activation of Nrf2 signaling in cortical astrocytes, whereas a protective antioxidant activity may be induced in other subtypes of astrocytes; and (ii) soy polyphenols and equol, but not daidzein, interfere with UPS function in primary cortical astrocytes.

**Figure 6.**
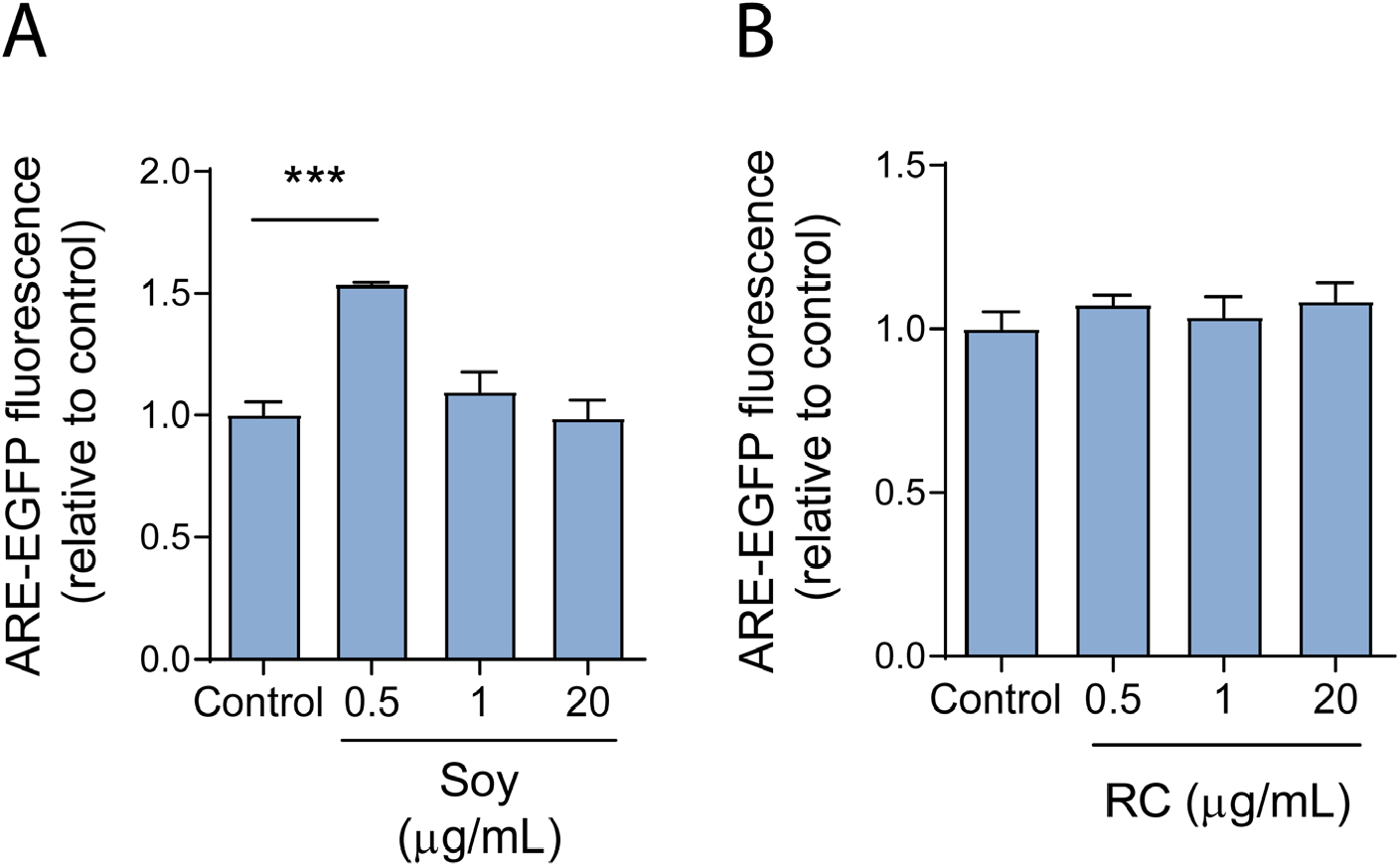
An extract prepared from soy but not red clover activates the Nrf2/ARE antioxidant response in hiPSC-derived astrocytes. (A,B) Human iPSC-derived astrocytes transduced with an ARE-EGFP reporter adenovirus for 48 h were incubated in the absence (‘control’) or presence of soy extract (A) or red clover extract (‘RC’) (B) for 24 h and imaged to determine the intracellular EGFP fluorescence intensity. The data are presented as the mean ± SEM; n = 3 (A) or n = 4 (B); ***p≤0.001, log transformation, one-way ANOVA with Dunnett’s multiple comparisons *post hoc* test.

### 3.6. Effects of red clover and soy extracts on rotenone-induced impairment of mitochondrial respiration

Several lines of evidence suggest that neurotoxicity elicited by rotenone and aSyn results from various pathogenic events including disruption of mitochondrial respiration and oxidative damage^9, 71, 94–96^. Accordingly, we hypothesized that the red clover and soy extracts might protect dopaminergic neurons against toxicity elicited by rotenone or aSyn by alleviating mitochondrial dysfunction. To address this hypothesis, we developed an assay aimed at measuring O_2_ consumption in human SH-SY5Y neuroblastoma cells challenged with rotenone in the absence or presence of extract prior to performing the assay. To ensure that the cells relied on oxidative phosphorylation as a primary source of energy production, the cultures were maintained in media containing galactose instead of glucose^40, 97, 98^. Cells exposed to rotenone exhibited a ~40% decrease in the rate of O_2_ consumption compared to untreated cells, and this decrease was largely reversed in cells pre-treated with extracts prepared from red clover (1 μg/mL) or soy (1 μg/mL), or with daidzein (50 nM) (Fig. 7A). These results suggested that the two extracts and daidzein interfered with rotenone-induced mitochondrial functional deficits.

**Figure 7.**
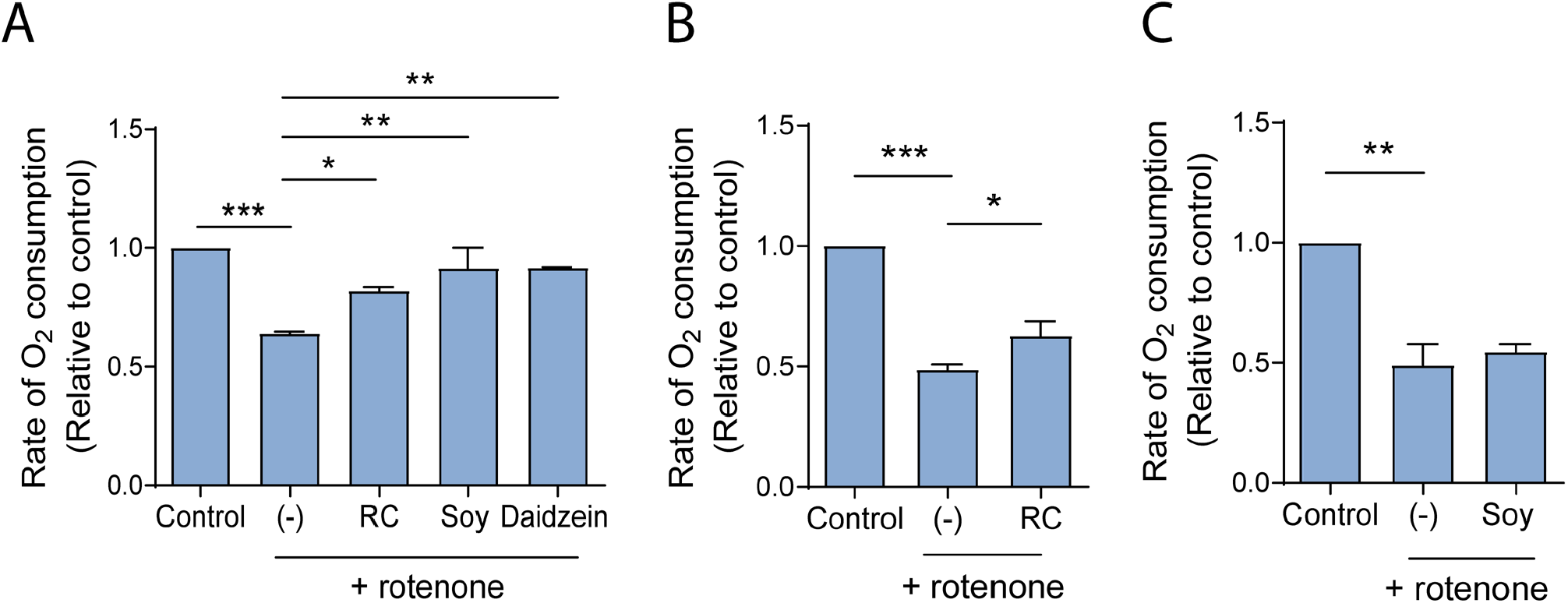
Isoflavone-rich extracts prevent rotenone-induced loss of mitochondrial function. (A) Galactose-conditioned SH-SY5Y cells were incubated in the absence or presence of a red clover extract (‘RC’) (1 μg/mL), soy extract (1 μg/mL), or daidzein (50 nM) for 19 h and then exposed to rotenone (30 nM) with or without extract or compound for 5 h. Control cells were incubated in the absence of rotenone, extract, or compound for 24 h. O_2_ consumption rates were measured in a suspension of cells with a Clark type oxygen electrode attached to a voltmeter. (B, C) O_2_ consumption rates were measured in a suspension of galactose-conditioned SH-SY5Y cells prepared in buffer containing rotenone (50 nM) with or without red clover extract (1 μg/mL) (B) or soy extract (1 μg/mL) (C). Control cells were analyzed in the absence of rotenone or extract. The data in (A-C) are presented as the mean ± SEM; n = 2 or 3; *p≤0.05, **p≤0.01, ***p≤0.001, log transformation, one-way ANOVA with Dunnett’s multiple comparisons *post hoc* test.

A number of polyphenols have been reported to compete with rotenone for ubiquinone binding sites in complex I of the electron transport chain^99^. Thus, we hypothesized that the red clover and soy extracts stimulate mitochondrial respiration by displacing rotenone from its binding site in complex I. To address this hypothesis, we developed a competition assay aimed at monitoring the effects of polyphenols on rotenone-mediated deficits in mitochondrial O_2_ consumption. Galactose-conditioned SH-SY5Y cells incubated in the presence of rotenone in the O_2_ consumption assay buffer exhibited a decrease in the rate of O_2_ consumption compared to untreated cells, and this inhibitory effect was partially rescued upon adding the red clover extract, but not the soy extract (Fig. 7B,C). These results suggest that the red clover extract, but not the soy extract, preserves mitochondrial function at least in part by displacing rotenone from its binding site in complex I.

### 3.7. Effects of a soy extract on 6-OHDA-induced motor dysfunction

To investigate the potential neuroprotective activity of soy isoflavones in an *in vivo* PD model, we tested a soy extract for the ability to alleviate motor deficits in 6-OHDA-treated rats, a classic neurotoxin model of PD. Unilateral, stereotaxic injection of 6-OHDA in the MFB or striatum results in a progressive phenotype involving dopaminergic cell death and depletion of striatal dopamine on the lesioned side^100–102^. The resulting imbalance in striatal dopamine causes pronounced motor deficits. For these experiments, we used another soy extract, Novasoy 400, which is enriched in genistin, daidzein, and glycitin. Rats received daily ip injections of extract or control vehicle prior to and after lesioning with 6-OHDA in the right MFB (Fig. 8A). Strikingly, rats injected with soy extract exhibited less pronounced motor defects in the open field test compared to rats treated with vehicle (Fig. 8B). Two-way ANOVA revealed a significant effect of time (p≤0.0001), treatment (p≤0.0001), and time x treatment (p≤0.0001). Lesioned animals treated with soy extract showed an increase in the percentage area covered at each time point compared to lesioned animals treated with vehicle, with a maximum increase of ~2-fold on day 10 (i.e. 9 days after 6-OHDA infusion). These results indicate that soy polyphenols alleviate motor dysfunction exhibited by rats exposed to 6-OHDA, potentially by interfering with nigrostriatal degeneration in these animals.

**Figure 8.**
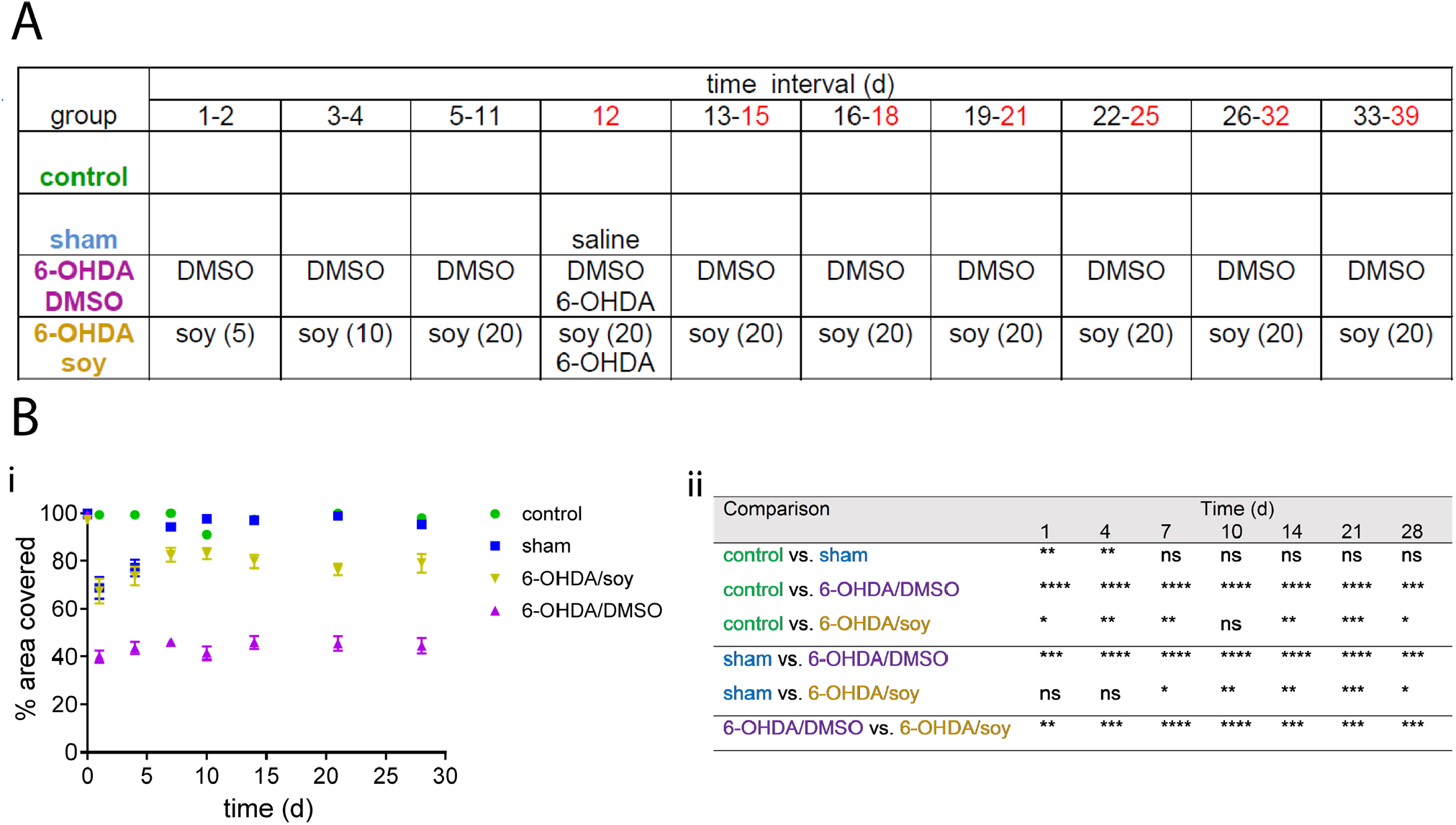
A soy extract alleviates motor deficits exhibited by 6-OHDA-lesioned rats. (A) Table outlining the experimental design. Numbers in the ‘6-OHDA/soy’ row refer to the amount of soy extract injected (mg/kg). Times shown in red at the top refer to days when motor tests were performed after 6-OHDA lesioning (days 12 to 39 of the overall study correspond to days 1 to 28 of open field testing shown in panel B). (Bi) Graph showing mean % area covered in the open field test at different times. (B") *P* values for open field data in panel Bi. The data are presented as the mean ± SEM; n = 3 (control) or n = 7 (sham, 6-OHDA/DMSO, 6-OHDA/soy); *p≤0.05, **p≤0.01, ***p≤0.001, ****p≤0.0001, log transformation, two-way ANOVA with Tukey’s multiple comparisons *post hoc* test (ns, non-significant). There were no significant differences among any of the groups at 0 d (i.e. day 11 of the overall study, 1 d prior to the injections).

## 4. Discussion

### 4.1. Isoflavone-rich extracts and individual isoflavones protect against neurotoxicity elicited by rotenone and A53T aSyn

The objective of this study was to examine the neuroprotective capacity and mechanisms of action of two isoflavone-rich extracts, prepared from red clover and soy, and the individual isoflavones daidzein and equol, a daidzein metabolite. We found that all of the extracts and isoflavones tested alleviated (or showed a trend towards alleviating) rotenone- and A53T-induced neurotoxicity in a primary midbrain culture model of PD (Figs. 1 and 5). These findings are consistent with previous data showing a rescue of dopaminergic cell loss in 6-OHDA-lesioned rats supplemented with a red clover diet^103^, and the ability of individual isoflavones to attenuate motor and non-motor symptoms induced by exposure to the toxins MPTP and 6-OHDA^34, 35^. Because the red clover and soy extracts examined in this study consist of a complex mixture of isoflavones, we infer that synergistic interactions involving multiple isoflavones could potentially play a role in the extracts’ neuroprotective activities^104–106^. At the same time, our observation that daidzein and equol interfered with toxicity elicited by rotenone or A53T aSyn indicates that individual isoflavones can also achieve neuroprotective effects in PD models. Interestingly, equol is a metabolite of daidzein produced by the microbiota of some but not all individuals. It has been suggested that the nature of consumed isoflavones (glycosides versus aglycones) could influence the production of equol in humans^107^, thus highlighting the importance of the plant material source of isoflavones with respect to their biological activity. Equol has shown health-promoting effects in models of stroke-like injury, osteoporosis, cardiovascular disease, and menopause^108–111^ and was recently found to alleviate toxicity elicited by the PD-related toxins MPP^+^ and 6-OHDA in SH-SY5Y cells^112^. Here, we show that equol can have neuroprotective effects in PD models at concentrations similar to plasma levels of the compound reported for equol producers^113^. These findings set the stage for future studies of neuroprotective pathways activated by daidzein and equol, with a view towards obtaining insights relevant to the development of personalized therapies accounting for microbiome differences among PD patients.

A number of isoflavones or their metabolites have been shown to penetrate the BBB in Sprague-Dawley rats^114, 115^, and equol was found to be highly permeable in an artificial BBB permeability assay^116^. Accordingly, we infer that the isoflavones and isoflavone-rich extracts examined in this study could potentially be of clinical benefit by reducing PD risk or slowing neurodegeneration in the brains of patients. The fact that daidzein and equol interfered with dopaminergic cell death at nanomolar concentrations in our primary midbrain culture model (Fig. 5C-E and G) implies that even a modest accumulation of these compounds in the brain could lead to neuroprotective effects in humans. Importantly, the ability of the extracts and individual isoflavones to mitigate neurotoxicity elicited by rotenone or A53T aSyn implies that these agents could potentially interfere with neurodegeneration in individuals with elevated PD risk triggered by a range of PD-related stresses. Consistent with this idea, we have obtained evidence from an ethnopharmacological study that a red clover extract is used as a form of traditional medicine by the Pikuni-Blackfeet Native Americans to treat PD-related symptoms^27^, and additional epidemiological data suggest that the consumption of isoflavones is associated with a reduced risk of neurodegenerative diseases^28, 29^.

### 4.2. A red clover extract activates the Nrf2/ARE antioxidant pathway and interferes with UPS function in cortical astrocytes

A key result of this study was our finding that a red clover extract, but not a soy extract or the individual isoflavones daidzein and equol, activated Nrf2 signaling in primary cortical astrocytes. We infer that Nrf2 underwent nuclear translocation in cells treated with extract based on our EGFP imaging data (in the case of cells transduced with ARE-EGFP reporter virus) and qRT-PCR data, both of which showed evidence of the transcription of Nrf2-regulated genes. In addition, Nrf2 nuclear translocation was previously shown to occur in parallel with the activation of an ARE-luciferase reporter closely related to the ARE-EGFP reporter used here^117^.

There is conflicting evidence in the literature about the ability of isoflavones to activate the Nrf2/ARE pathway and induce the expression of Nrf2-regulated genes. For example, two isoflavone metabolites, tectorigenin and glycitein, were found to up-regulate HO1 and NAD(P)H Quinone Dehydrogenase 1 (NQO1) expression in a Nrf2-dependent manner in cortical astrocytes^118^. Daidzein was shown to up-regulate HO1 in smooth muscle-derived cells but failed to increase quinone reductase (QR) activity in colon cancer cells^119^. In another study, daidzein and genistein were found to increase QR activity in different tissues of mice fed a daidzein- or genistein-enriched diet^120^. Interestingly, the latter study revealed variable tissue- and sex-dependent effects of isoflavone administration on glutathione-S-transferase activity^120^. These reported differences across various tissues suggest a possible cell type-specific response to isoflavones that could explain the lack of Nrf2 activation by the soy extract and individual isoflavones in cortical astrocytes (Supplementary Fig. 4).

This is the first report of UPS inhibition by a red clover extract (Fig. 4A). A number of polyphenols, including isoflavones, have been shown by our group (Supplementary Fig. 5) and others to interfere with UPS activity^79, 81, 121–123^. Genistein, a soy isoflavone, was reported to inhibit the chymotrypsin-like activity of the proteasome in a cancer cell model, likely by interacting with the proteasomal β5 subunit^123^. Our observation that a red clover extract interfered with GFPu degradation implies that polyphenols in this extract could induce activation of the Nrf2/ARE pathway by inhibiting UPS function. The concentration at which the extract induced a significant increase in the GFPu signal is higher than the concentration used for Nrf2 activation, implying that the sensitivity of the GFPu reporter assay is less than that of the ARE-EGFP assay, and, therefore, a low level of proteasome inhibition may be sufficient to induce Nrf2 signaling but not sufficient to cause a measurable amount of GFPu accumulation. In contrast, red clover polyphenols apparently do not activate Nrf2-mediated transcription via a mechanism involving redox cycling reactions leading to Keap1 oxidation, given that the antioxidant molecule NAC had no impact on the ability of the red clover extract to activate Nrf2 signaling in cortical astrocytes (Fig. 3)^48, 60, 124^. Finally, our observation that the red clover extract induced a modest up-regulation of Nrf2 mRNA levels (Supplementary Fig. 3) suggests that this is an additional mechanism by which red clover polyphenols could activate Nrf2 signaling, consistent with similar effects reported for quercetin and resveratrol^88, 89^.

### 4.3. Activation of astrocytic Nrf2 signaling by a red clover extract depends on the brain region from which the astrocytes are prepared

Another important outcome of our study was the finding that astrocytic Nrf2-mediated transcription was activated by a red clover extract in astrocytes prepared from rat cortex (Fig. 2C,D), but not in astrocytes prepared from rat midbrain or in mixed midbrain cultures (Supplementary Fig. 1). These observations are consistent with previous data showing that astrocytes from different brain regions differ in terms of their molecular profiles and their responses to different stimuli^125–132^. For example, midbrain and cortical astrocytes were found to express different levels of glial glutamate transporter-1 (GLT-1) in response to corticosteroid stimulation^133^. The data presented here support previous findings relating to the heterogeneity of glial cells^125, 130, 134^ by showing that astrocytes prepared from different brain regions respond differently to inducers of the Nrf2-mediated antioxidant response. Another explanation could be that red clover polyphenols undergo metabolic reactions that result in a loss of the extract’s ability to modulate the Nrf2/ARE pathway in midbrain but not cortical astrocytes. Our data could also be explained by differences in the timing of Nrf2 activation among different types of astrocytes, although our observation that all of the astrocytic cultures examined here showed similar degrees of Nrf2-dependent signaling after treatment with curcumin for 24 h (Supplementary Fig. 2B) indicates that at least some forms of polyphenol-mediated Nrf2 activation can occur with similar kinetics in astrocytes from different brain regions. This observation also suggests that (i) curcumin and the red clover extract activate Nrf2 signaling via different mechanisms; or (ii) curcumin is resistant to metabolic reactions that interfere with Nrf2 activation by red clover polyphenols in mixed midbrain cultures. Finally, we note that the activation of our ARE-EGFP reporter derived from the mouse HO-1 gene presumably only reflects the expression of a subset of all 200+ Nrf2 targets. Accordingly, additional Nrf2-dependent transcriptional events could occur even in isoflavone-treated astrocytes that show no fluorescent signal in our reporter assay.

Collectively, our results suggest that the ability of isoflavone-rich extracts to activate astrocytic Nrf2 signaling varies in astrocytes and mixed cultures prepared from different brain regions. Our observation that the red clover and soy extracts exhibited opposite patterns of Nrf2 activation in rat cortical astrocytes versus human iPSC-derived astrocytes (Figs. 2 and 6 and Supplementary Fig. 4A) further highlights the point that isoflavone-rich extracts induce up-regulation of Nrf2-mediated transcription in a manner that varies across different types of astrocytic cultures. Uncovering mechanisms underlying these brain region- and cell culture-specific responses has the potential to advance our understanding of the role of astrocytes in neuroprotection and enhance the impact of medicines designed to target the Nrf2/ARE pathway.

Lastly, because the red clover extract failed to activate Nrf2 signaling monitored using the ARE-EGFP reporter in primary midbrain cultures, we infer that the extract’s protective effects against toxicity elicited by rotenone or A53T aSyn did not involve the up-regulation of astrocytic, Nrf2-mediated transcriptional events reflected by the reporter (e.g. HO-1 gene transcription) under our experimental conditions. Nevertheless, the extracts could potentially alleviate dopaminergic cell death in midbrain cultures exposed to PD-related insults via Nrf2 signaling mechanisms not reflected by our ARE-EGFP reporter, as discussed above.

### 4.4. Isoflavone-rich extracts rescue mitochondrial dysfunction

O_2_ consumption experiments revealed that the red clover and soy extracts alleviated rotenone-induced deficits in mitochondrial electron transport in SH-SY5Y neuroblastoma cells (Fig. 7A). These findings are consistent with previous data showing that soy isoflavones mitigated brain mitochondrial oxidative stress triggered by the amyloid-β peptide in rat models of AD^135, 136^. Based on evidence that the neurotoxic effects of rotenone exposure and aSyn over-expression are related to mitochondrial functional deficits^9, 71, 94–96^, we infer that a rescue of these deficits by the red clover and soy extracts could play a key role in the extracts’ ability to protect against these insults. Our finding that the red clover extract, but not the soy extract, preserved mitochondrial function at least in part by displacing rotenone from its binding site in complex I (Fig. 7B,C) suggests that red clover polyphenols can protect against rotenone neurotoxicity by interfering with the toxicant’s ability to disrupt mitochondrial electron transport. Previous studies revealed that isoflavones such as genistein interact with the electron transport chain^137^, and genistein and daidzein were found to associate with the F0F1-ATPase/ATP synthase^138^ and/or modulate NADH:ubiquinone oxidoreductase activity^139^. Our observation that the soy extract failed to displace rotenone from its complex I binding site (Fig. 7C) implies that only specific isoflavones rescue mitochondrial functional deficits via interactions with electron transport chain components. In addition to engaging in such interactions, isoflavones in both the red clover and soy extracts could ameliorate mitochondrial dysfunction via other mechanisms, including potentially up-regulating the transcriptional co-activator PGC1-alpha, a master regulator of mitochondrial biogenesis^140^. Moreover, isoflavones could enhance mitochondrial function and quality control by promoting the activity of the E3 ubiquitin ligase parkin^141, 142^, a protein known to be impaired in cells exposed to PD-related insults including extracellular aSyn oligomers^143, 144^. Additional insight into mitochondrial mechanisms underlying isoflavone-mediated neuroprotection could be obtained via (i) biochemical analyses of neuronal lysates to measure the activities of mitochondrial enzymes such as those outlined above; and (ii) TEM analyses of intact (fixed) neurons to assess the effects of isoflavones on rotenone-dependent changes in mitochondrial morphology^145, 146^.

### 4.5. An isoflavone-enriched soy extract protects against 6-OHDA toxicity *in vivo*

Treatment of 6-OHDA-lesioned rats with Novasoy 400, a 40% (w/v) isoflavone standardized soy extract enriched in genistin, daidzein, and glycitin, led to an amelioration of 6-OHDA-induced motor dysfunction (Fig. 8). These findings suggest that soy isoflavones can mitigate nigrostriatal degeneration induced by 6-OHDA, potentially by rescuing mitochondrial dysfunction and activating antioxidant pathways in astrocytes, as seen in our cellular models of PD. Consistent with this idea, isoflavones such as genistein and equol have been shown to cross the BBB to reach target cell types^114, 116^. The ability of a standardized soy extract to improve motor function in 6-OHDA-treated rats is in agreement with other published studies showing neuroprotective activities of individual soy isoflavones in animal models of PD. For example, the abundant soy isoflavone genistein was shown to increase the survival of neurons in the *substantia nigra* of rodents exposed to 6-OHDA^36^ or MPTP^34^, and the daidzein metabolite equol protected *Caenorhabditis elegans* against toxicity elicited by MPP^+^^112^. Future studies will be focused on examining neuroprotective effects of Novasoy 400 via immunohistochemical analysis of brain sections from rats exposed to 6-OHDA and other PD-related insults.

## 5. Conclusion

In conclusion, isoflavone-rich red clover and soy extracts, and the individual isoflavones daidzein and equol, were found to alleviate neurotoxicity elicited by insults linked epidemiologically or genetically to PD. Equol showed potent neuroprotective activity at levels similar to those in the plasma of equol-producers, suggesting that there may be opportunities for the development of individualized therapies targeting patients with particular gut microbiome profiles. Although both the red clover and soy extracts interfered with neurotoxic effects of PD-related insults, they differed in their abilities to modulate astrocytic Nrf2 signaling and ameliorate mitochondrial dysfunction, highlighting the important role of the extracts’ polyphenolic composition and potential synergies among their constituents in determining their neuroprotective mechanisms. Based on evidence that a number of isoflavones or their metabolites can penetrate the BBB^114, 115^, our findings suggest that the extracts and isoflavones examined in this study have the potential to lower the risk of PD or slow disease progression. Studies in humans revealed that consuming 60 or 100 mg total isoflavone equivalent per day for 10 to 12 weeks resulted in improved cognitive function^31, 147^, suggesting that isoflavone supplementation is a viable strategy to promote brain health. An important area of future research will be to determine whether isoflavones interact with existing PD medications, leading to adverse effects in patients. Although animal studies have shown evidence of toxicity resulting from long-term or high-dose intake of isoflavones (e.g. 150 mg/kg body weight of genistein)^148^, a phase I clinical trial reported that a high daily dose of isoflavones was well-tolerated in healthy women^149^. A more complete understanding of the risks and benefits of isoflavone administration in humans would advance efforts to develop safe and efficacious dietary interventions for PD patients.

## Author Contributions

**A.d.R.J.**: conceptualization, formal analysis, investigation, methodology, visualization, writing – original draft, writing – review & editing. **A.A.**: formal analysis, investigation, methodology, visualization. **M.A.T.**: formal analysis, investigation, methodology, visualization, writing – review & editing. **S.Y.M.**: formal analysis, investigation. **M.T.**: formal analysis, investigation, methodology, visualization. **M.H.G.**: formal analysis, visualization, writing – review & editing. **Q.- L.W.**: resources, writing – review & editing. **J.E.S.**: resources, writing – review & editing. **G.P.M.**: formal analysis. **M.A.L.**: resources, supervision, writing – review & editing. **R.S.**: resources, supervision, writing – review & editing. **J.-C.R.**: conceptualization, funding acquisition, project administration, resources, supervision, visualization, writing – review & editing.

## Acknowledgements

This work was supported by NIH grants R21 AG039718 and R03 DA027111 (J.-C. R), a Pilot Grant from the Purdue-UAB Botanicals Research Center (NIH P50 AT000477-06), grants from the Branfman Family Foundation and Showalter Trust (J.-C.R.), a Soy Health Research Program Incentive Award from the United Soybean Board (J.-C.R.), a fellowship from the Botany in Action program, Phipps Botanical Garden, Pittsburgh (A.d.R.J.), and a fellowship from the Purdue Research Foundation (A.d.R.J.). The research described herein was conducted in a facility constructed with support from Research Facilities Improvement Program Grants number C06-14499 and C06-15480 from the National Center for Research Resources of the NIH. The authors would like to thank the members of the laboratory for important discussions and feedback.

---

**Supplementary Figure 1.**
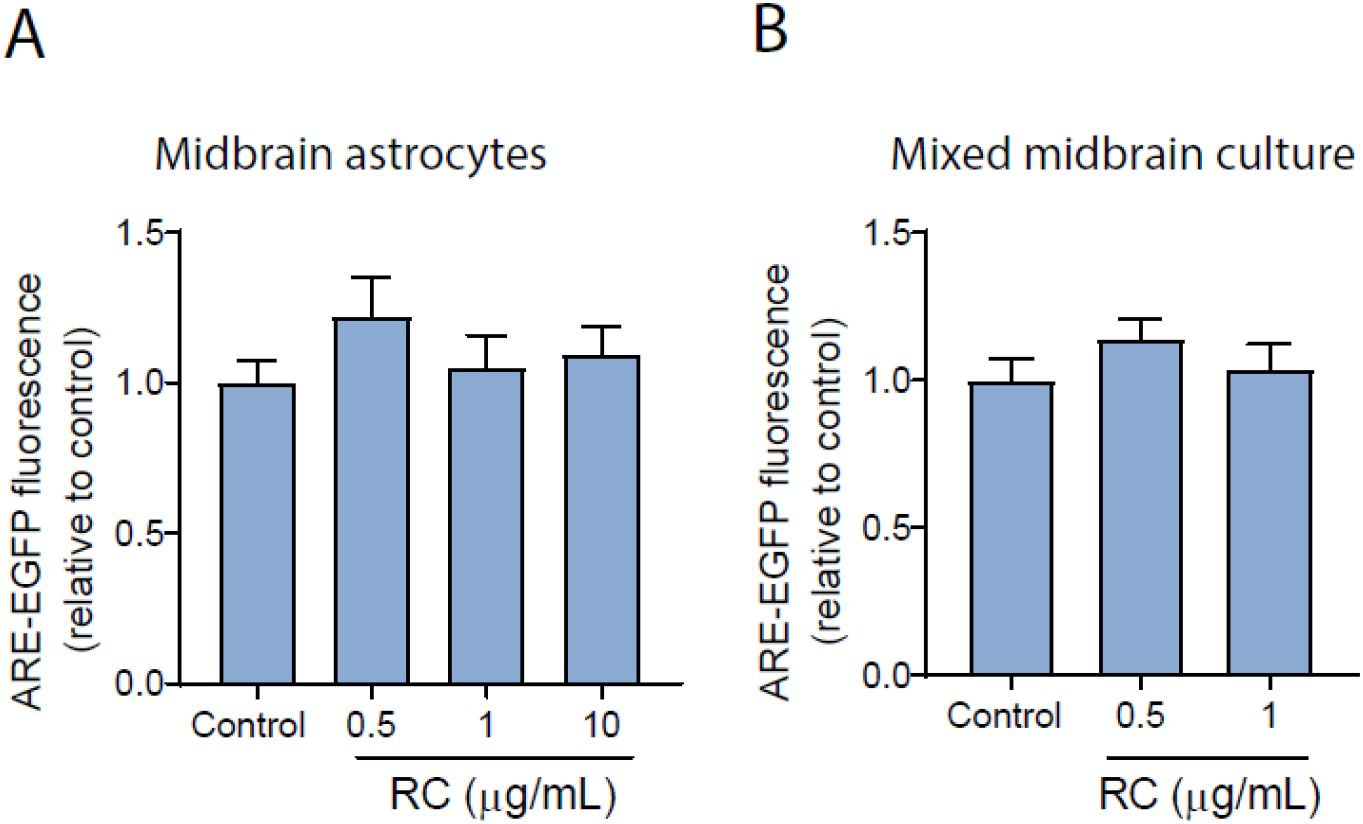
A red clover extract fails to activate Nrf2-mediated transcription in midbrain astrocytes. Primary midbrain astrocytes (A) or mixed midbrain cultures (B) transduced with an ARE-EGFP reporter adenovirus for 48 h were incubated in the absence (‘control’) or presence of red clover extract (‘RC’) for 24 h and imaged to determine the intracellular EGFP fluorescence intensity. The data are presented as the mean ± SEM; n = 6 (A) or n = 3 (B).

**Supplementary Figure 2.**
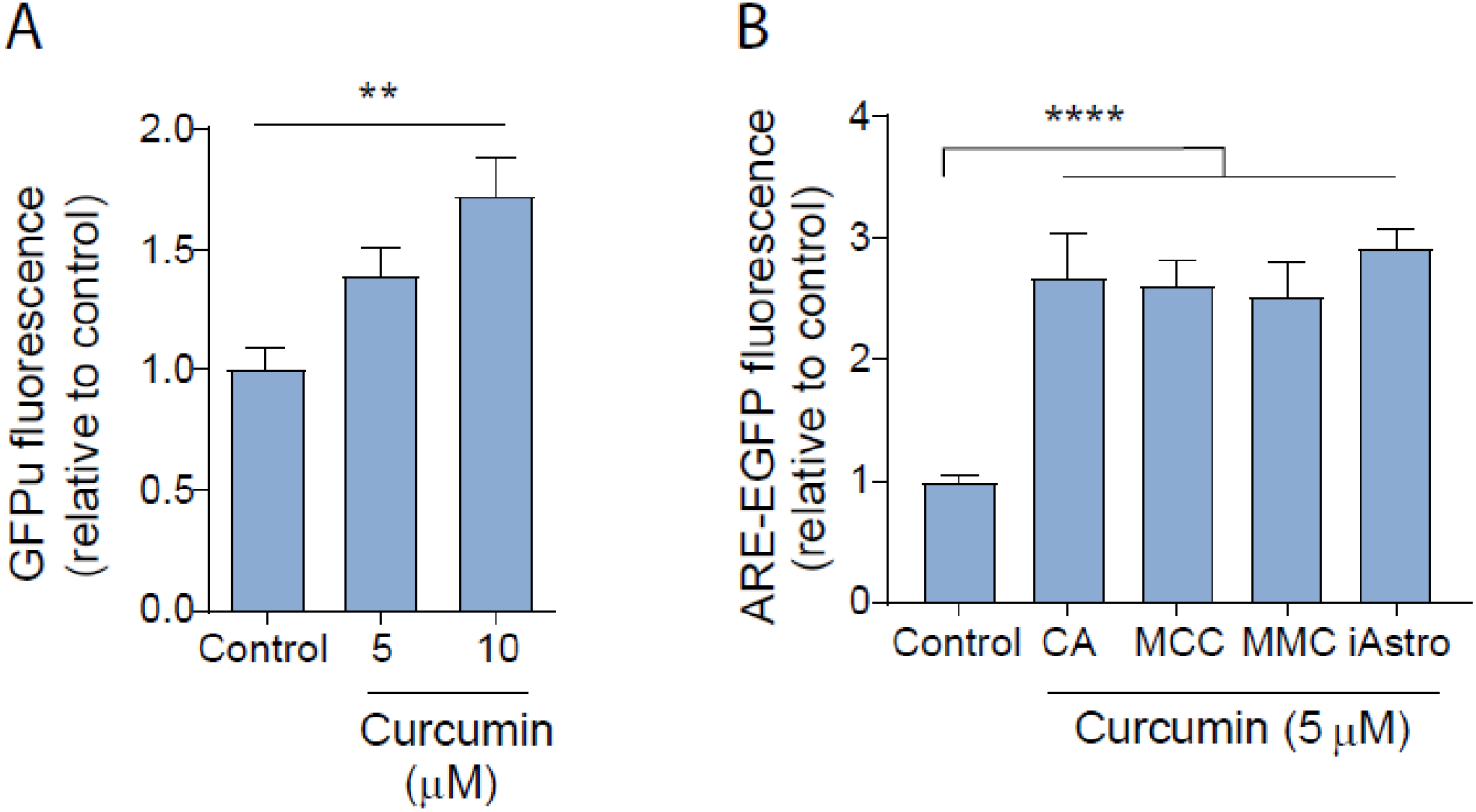
Curcumin inhibits the UPS and activates Nrf2-mediated transcription. (A) Primary cortical astrocytes transduced with a reporter adenovirus encoding the UPS substrate GFPu for 48 h were incubated in the absence (‘control’) or presence of curcumin for 24 h and imaged to determine the intracellular GFP fluorescence intensity. (B) Primary cortical astrocytes (CA), mixed cortical cultures (MCC), mixed midbrain cultures (MMC), or iPSC-derived astrocytes (iAstro) were transduced with an ARE-EGFP reporter adenovirus for 48 h and incubated in the absence (‘control’) or presence of curcumin for 24 h. The cells were imaged to determine the intracellular EGFP fluorescence intensity. The data in (A) and (B) are presented as the mean ± SEM; n = 3; **p≤0.01, ****p≤0.0001, log transformation, one-way ANOVA with Dunnett’s multiple comparisons *post hoc* test. The control value in (B) was determined by pooling the data obtained for the different types of cell cultures incubated without curcumin (n = 12 in total).

**Supplementary Figure 3.**
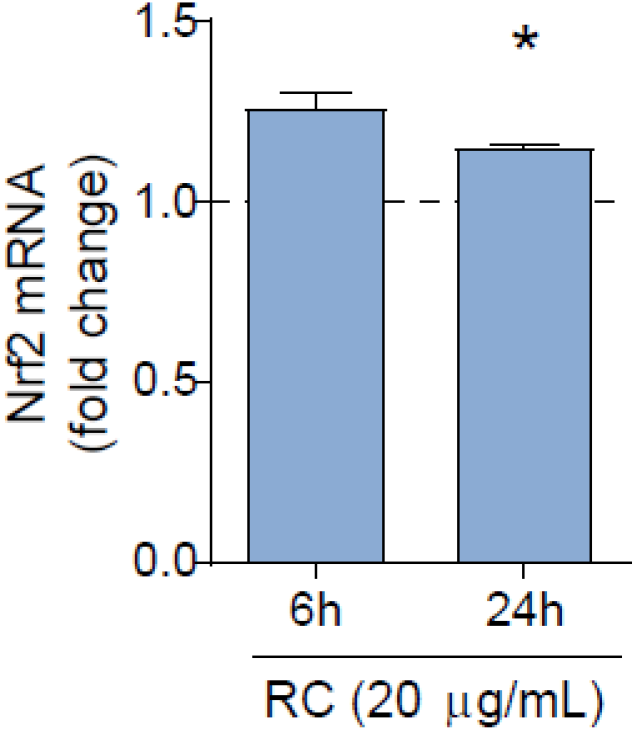
A red clover extract induces a modest upregulation of Nrf2 expression. Primary cortical astrocytes were incubated in the absence or presence of a red clover extract (‘RC’) for 6 h or 24 h, and Nrf2 mRNA levels were measured by qRT-PCR. The data are presented as the mean ± SEM; n = 2; *p≤0.05 versus a predicted ratio of 1, log transformation followed by one-sample t-test.

**Supplementary Figure 4.**
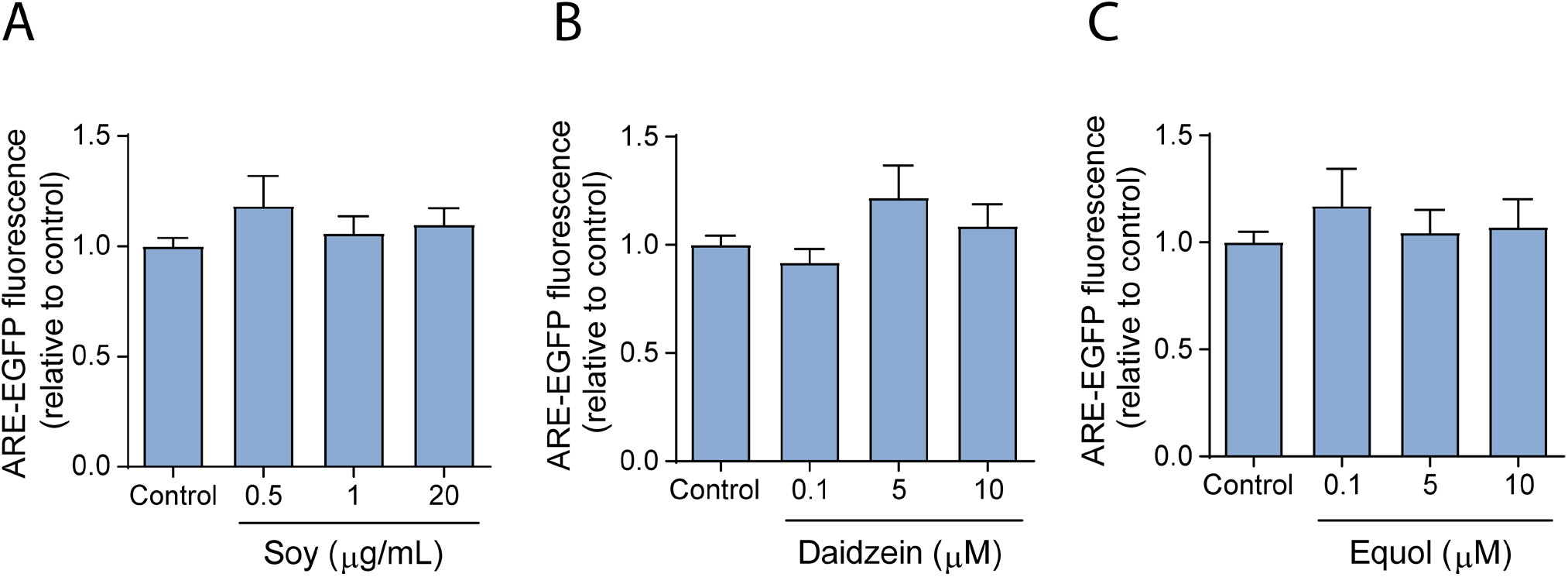
A soy extract and soy isoflavones fail to activate Nrf2-mediated transcription in cortical astrocytes. Primary cortical astrocytes transduced with an ARE-EGFP reporter adenovirus for 48 h were incubated in the absence (‘control’) or presence of soy extract (A), daidzein (B), or equol (C) for 24 h and imaged to determine the intracellular EGFP fluorescence intensity. The data are presented as the mean ± SEM; n = 9 (A); n = 7 (B); n = 6 (C).

**Supplementary Figure 5.**
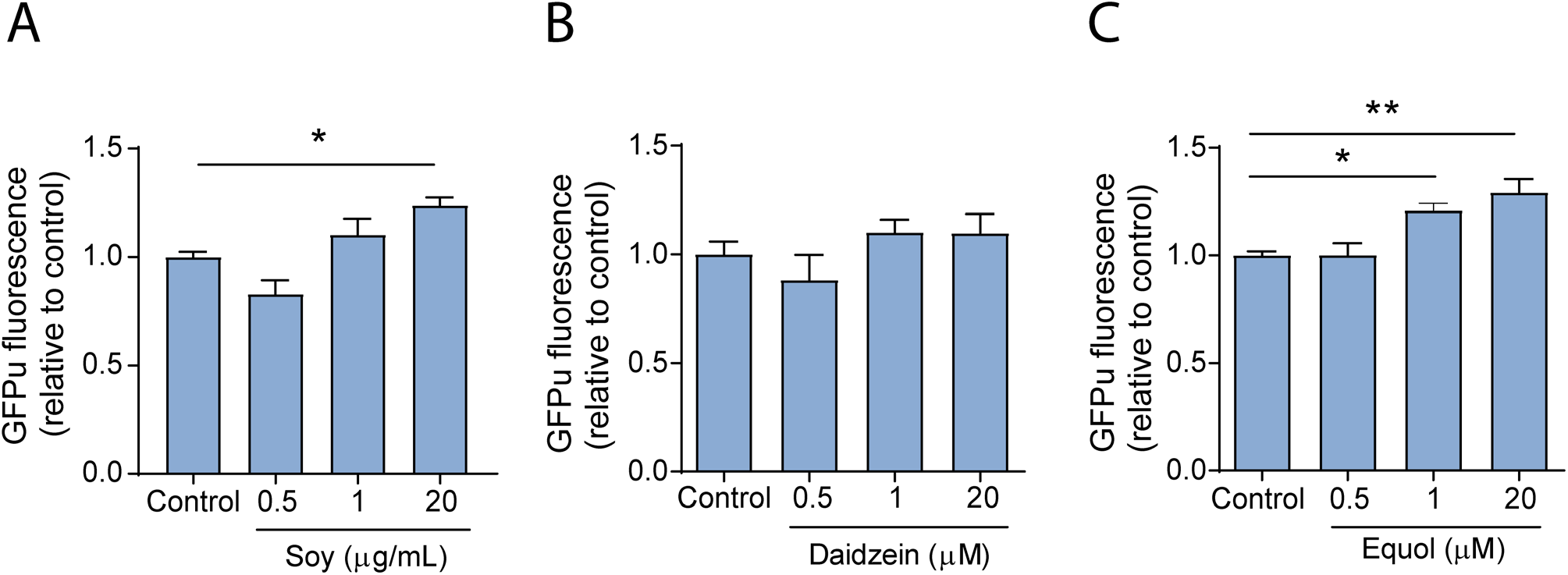
A soy extract and equol (but not daidzein) inhibit the UPS. Primary cortical astrocytes transduced with a reporter adenovirus encoding the UPS substrate GFPu for 48 h were incubated in the absence (‘control’) or presence of soy extract (A), daidzein (B), or equol (C) for 24 h and imaged to determine the intracellular GFP fluorescence intensity. The data are presented as the mean ± SEM; n = 4 (A), n = 7 (B), n = 3 (C); *p≤0.05, **p≤0.01, log transformation, one-way ANOVA with Dunnett’s multiple comparisons *post hoc* test.

